# Convergent Insulin and TGF-β signalling drives cancer cachexia by promoting aberrant fatbody ECM accumulation in a *Drosophila* tumour model

**DOI:** 10.1101/2023.06.10.544444

**Authors:** Daniel Bakopoulos, Sofia Golenkina, Callum Dark, Elizabeth L Christie, Besaiz J. Sánchez-Sánchez, Brian M. Stramer, Louise Y Cheng

**Affiliations:** Peter MacCallum Cancer Centre, Melbourne, VIC 3000, Australia; Sir Peter MacCallum Department of Oncology, The University of Melbourne, VIC 3010, Australia; Department of Anatomy and Physiology, The University of Melbourne, VIC 3010, Australia; Kings College London, Randall Centre for Cell & Molecular Biophysics, London SE1 1UL, United Kingdom

**Keywords:** *Drosophila*, cachexia, insulin, TGF-β, Sog, ECM, SPARC

## Abstract

Cancer cachexia is a wasting disease suffered by advanced stage cancer patients and ultimately causes ∼30% of cancer mortalities. Clinical observations have shown that extracellular matrix (ECM) remodelling which leads to fibrosis in the adipose tissue is a key feature of cancer cachexia. However, the molecular regulators of adipose ECM remodelling are not known and how this leads to muscle wasting is unclear. In this study, using a *Drosophila* cachexia model, we found that in the adipose tissue of both wildtype and tumour bearing animals, insulin and TGF-β signalling converge via a BMP antagonist *short gastrulation* (*sog*) to regulate ECM remodelling. In tumour bearing animals, the aberrant ECM accumulation in the fatbody, contributes towards muscle detachment by preventing ECM secretion and subsequently depleting muscles of fatbody-secreted ECM proteins. Strikingly, activation of insulin signalling, inhibition of TGF-β signalling, or modulation of ECM secretion via SPARC or Rab10 in the fatbody, was able to rescue tissue wasting in the presence of tumour. Together, our study highlights the importance of adipose ECM remodelling in the context of cancer cachexia.

## Introduction

Cachexia is a metabolic wasting syndrome characterised by the loss of adipose tissue and muscle. Cachexia occurs in approximately 80% of advanced-stage cancer patients and is ultimately responsible for approximately 30% of cancer mortalities. Despite the clinical significance of cachexia in cancer and diseases such as HIV or other chronic diseases, there is no gold standard for its treatment (Baracos et al., 2018). Cachexia is often resolved when the tumour can be resected in early disease, however, there are few strategies to treat cachexia in late stage or metastatic patients, where removal of the tumour is not an option. Therefore, understanding the mechanisms that drive wasting in peripheral tissues and how to target these deregulations can offer new therapeutic avenues to treat cachexia.

*Drosophila melanogaster* is emerging as an excellent model to identify tumour-secreted factors that drive cancer cachexia. Adult and larval tumour models that induce cachectic phenotypes have been established in *Drosophila*, and these models have revealed several different mechanisms by which tumours induce these cachectic phenotypes (Ding et al., 2021; Figueroa-Clarevega and Bilder, 2015; Hodgson et al., 2021; Khezri et al., 2021; Kwon et al., 2015; Lee et al., 2021; Lodge et al., 2021; Newton et al., 2020; Santabárbara-Ruiz and Léopold, 2021; Song et al., 2019). We recently demonstrated that eye imaginal disc tumours, caused by the expression of constitutively activate *Ras* (*Ras^V12^*) and an RNAi against the polarity protein disc-large 1 (*dlg^RNAi^*), secrete two cachectic factors: ImpL2 and Matrix metalloproteinase 1 (Mmp1) (Lodge et al., 2021). ImpL2 is a secreted protein that induces cachexia by binding to circulating Insulin-like peptides (Ilps) to prevent them from activating Insulin-like receptor (InR) in host tissues, thus causing host tissue wasting (Honegger et al., 2008). This mechanism is likely conserved, as Insulin Growth Factor Binding Protein-2 (IGFBP-2), which also antagonises insulin signalling, is correlated with muscle atrophy in pancreatic ductal adenocarcinoma (PDAC) patients (Dong et al., 2021). In addition, Mmp1 drives cachexia by increasing the amount of the Transforming growth factor-β (TGF-β) ligand Glass bottom boat (Gbb). Gbb subsequently induces an elevation of TGF-β signalling in the *Drosophila* adipose tissue (called the fatbody), to induce muscle detachment. Together, our findings highlighted the importance of fatbody signalling in promoting muscle degradation during cachexia. However, what occurs downstream of these signals to drive fatbody and muscle disruption remains unclear.

In this study, we demonstrate that tumour derived ImpL2 and Gbb facilitate muscle degradation during cachexia via two main mechanisms. Firstly, ImpL2-mediates a reduction in fatbody insulin signalling, which in turn activates fatbody TGF-β signalling by upregulating *short gastrulation* (*sog*), a TGF-β antagonist. As Gbb also activates fatbody TGF-β signalling, our findings reveal that ImpL2 and Gbb converge on this activation. We further show that fatbody TGF-β signalling activation subsequently causes an aberrant upregulation of ECM proteins in the fatbody. This deposition of disorganized ECM leads to fibrotic ECM accumulation. As muscle ECM proteins are mostly fatbody-derived (Dai et al., 2017; Dai et al., 2018), this in turn causes a reduction in muscle ECM to promote muscle detachment. Modulating SPARC, a collagen binding protein (Shahab et al., 2015) or Rab10, a regulator of basement membrane fibril formation (Isabella and Horne-Badovinac, 2016) can ameliorate ECM accumulation in the fatbody in tumour bearing animals, and in turn improve muscle integrity. Secondly, tumour secreted ImpL2 causes a reduction in muscle insulin signalling, which contributes towards reduced translation and increased muscle atrophy. Enhanced activation of insulin in the muscle can specifically improve muscle atrophy (by not ECM). Together, the two mechanisms attribute towards the muscle detachment phenotype we have observed. Therefore, our data demonstrate that in addition to targeting tumour secreted factors, modulating key signalling and ECM regulators in peripheral tissues could be an effective strategy to treat cachexia.

## Results

### Reduced insulin signalling promotes the activation of TGF-β signalling in the cachectic fatbody

In this study, we utilise two *Drosophila* larval tumour models to study cachexia. In the first model, the tumour is induced via the *GAL4-UAS* mediated overexpression of *Ras^V12^*and Disc Large (Dlg) RNAi in the eye (Figure 1A). Using this system, we can knockdown or overexpress candidate genes in the tumour. In the second model, the tumour is induced via the *QF2-QUAS* mediated overexpression of *Ras^V12^* and *scrib* RNAi (Figure 1B), allowing us to knockdown or overexpress genes of interest in the fatbody of tumour-bearing animals using drivers such as *R4-GAL4*. We previously showed that *Ras^V12^*, *dlg^RNAi^* eye imaginal disc tumours express elevated levels of the dIlp antagonist *ImpL2*, consequently leading to a reduction in fatbody insulin signalling (Lodge et al., 2021). In parallel, tumour secreted Gbb accounts for the upregulation of TGF-β signalling. Here, we show that tumours induced by the overexpression of *Ras^V12^* and an RNAi against the polarity protein scribble (*scrib^RNAi^)* using the QF/QUAS system, similarly caused a downregulation of insulin signalling (pAkt, Figure 1 C-E), and an upregulation of TGF-β signalling (pMad, Figure 1 F-H) in the fatbody.

**Figure 1:**
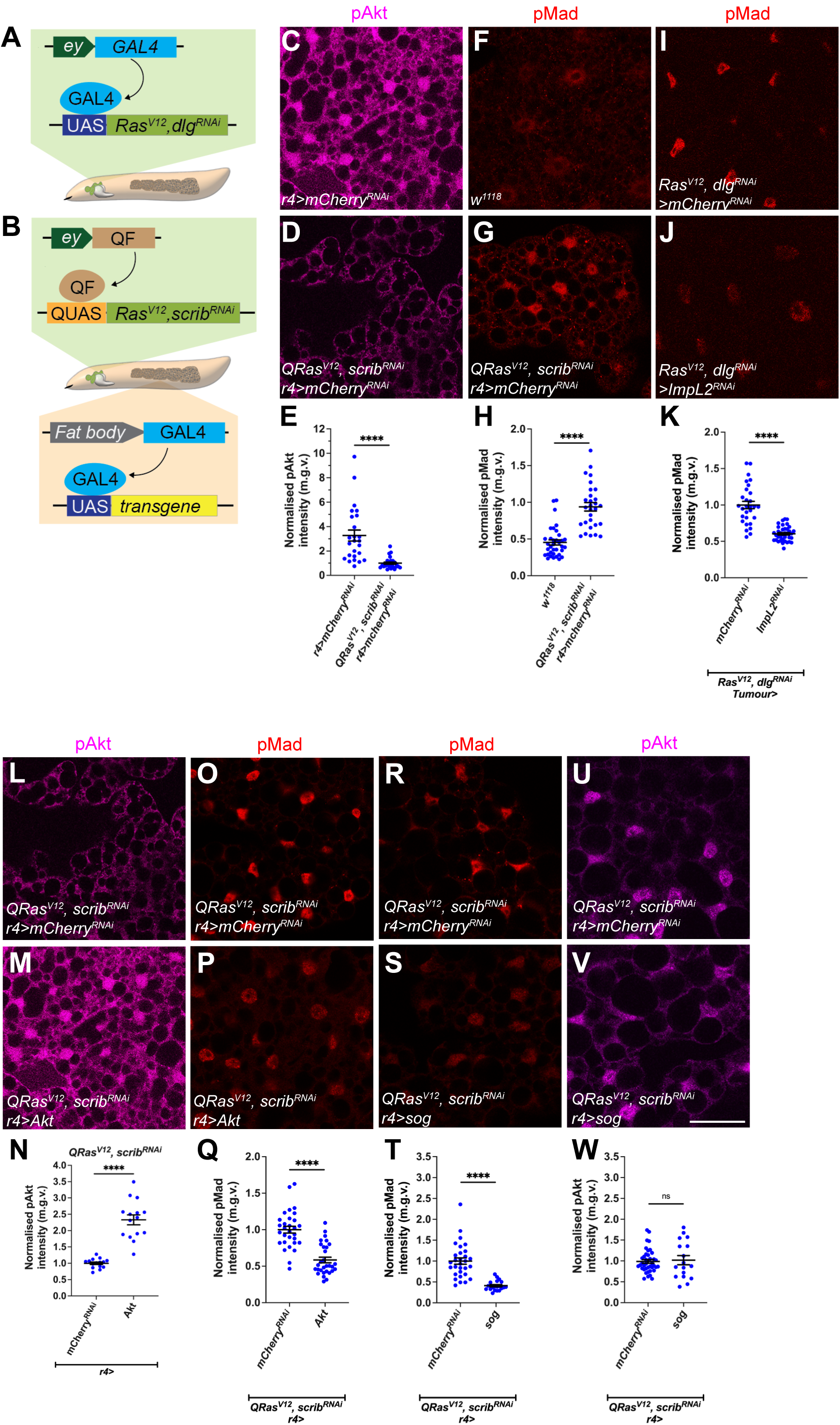
Insulin signalling in the fatbody negatively inhibits TGFß signalling during cachexia. (A-B) Schematic depicting the two tumour models utilized in this study. The GAL4-UAS induced *Ras^V12^;dlg^RNAi^*tumour, or the *QF2-QUAS* induced *Ras^V12^scrib^RNAi^*tumour. (C-D) Fatbody stained for pAkt from control (r4> *mCherry^RNAi^)* and *QRas^V12^scrib^RNAi^; r4>mCherry^RNAi^* tumour-bearing animals. (E) Quantification of normalized (to tumour) fatbody pAkt intensity in A-B*. r4>mCherry^RNAi^* (n=28), *QRas^V12^scrib^RNAi^; r4>mCherry^RNAi^* (n=25). (F-G) Fatbody stained for pMad from control (*w^1118^)* and *QRas^V12^scrib^RNAi^; r4>mCherry^RNAi^* tumour-bearing animals. (H) Quantification of normalized (to tumour) fatbody pMad intensity in F-G*. w^1118^* (n=32), *QRas^V12^scrib^RNAi^; r4>mCherry^RNAi^* (n=31). (I-J) Fatbody of *Ras^V12^dlg^RNAi^* tumour-bearing animals where *mCherry^RNAi^* or *ImpL2^RNAi^* was expressed in the tumour, stained for pMad. (K) Quantification of fatbody pMad intensity in I-J*. mCherry^RNAi^* (n=29), *ImpL2^RNAi^* (n=37). (L-M) Fatbody stained for pAkt of *QRas^V12^scrib^RNAi^* tumour-bearing animals where *mCherry^RNAi^* or *Akt* was expressed in the fat body (*r4-GAL4*). (N) Quantification of normalized fatbody pAkt intensity in L-M*. QRas^V12^scrib^RNAi^; r4> mCherry^RNAi^* (n=14), *QRas^V12^scrib^RNAi^; r4>Akt* (n=14). (O-P) Fatbody of *QRas^V12^scrib^RNAi^* tumour-bearing animals where *mCherry^RNAi^* or *Akt* was expressed in the fatbody (*r4-GAL4*), with TGF-ß signaling activation indicated by pMad staining. (Q) Quantification of normalized fatbody pMad intensity in O-P*. mCherry^RNAi^* (n=30), *Akt* (n=30). (R-S) Fatbody from animals expressing *mCherry^RNAi^* or *sog* under the control of *r4-GAL4* in *QRas^V12^scrib^RNAi^* tumour-bearing animals stained with pMad. (T) Quantification of normalized fatbody pMad intensity in R-S*. mCherry ^RNAi^* (n=30), *sog* (n=20). (U-V) Fatbody from animals expressing *mCherry^RNAi^* or *sog* under the control of *r4-GAL4* in *QRas^V12^scrib^RNAi^* tumour-bearing animals stained with pAkt. (W) Quantification of normalized fatbody pAkt intensity in U-V*. mCherry ^RNAi^* (n=40), *sog* (n=17). Scale bar is 50µm for fatbody pAkt and pMad staining

Tumour specific ImpL2 inhibition was sufficient to reduce fatbody pAkt levels (Lodge et al., 2021) and ameliorate tumour-induced muscle disruption (Figueroa-Clarevega and Bilder, 2015; Honegger et al., 2008; Kwon et al., 2015; Lee et al., 2021). Surprisingly, we found that the knockdown of ImpL2 in the tumour (*Ras^V12^ dlg^RNAi^*) caused a reduction in fatbody pMad staining (Figure 1 I-K), suggesting that there may be signalling crosstalk between the insulin/PI3K and TGF-β pathways in the fatbody. Furthermore, expression of *Akt* overexpression in the fatbody of tumour animals (*QRas^V12^, scrib^RNAi^*), caused an upregulation of pAkt (Figure 1L-N), but an unexpected downregulation of the TGF-β signalling readout pMad (Figure 1O-P), indicating that an upregulation of insulin/PI3K signalling maybe correlated with a downregulation of TGF-β signalling. However, conversely, TGF-β signalling does not seem to affect the activation of PI3K signalling pathway. Upon the expression of the TGF-β inhibitor *sog* (Biehs et al., 1996) in the fatbody (*r4-GAL4*) of tumour bearing animals (*QRas^V12^, scrib^RNAi^*); we observed a significant reduction in fatbody pMad levels (Figure 1R-T) but little effect on pAkt levels (Figure 1U-W).

### Enhancing insulin signalling in the fatbody improves muscle integrity in cachectic animals

Next, we tested whether fatbody-specific changes in Insulin signalling influences muscle breakdown during cachexia. Similar to *GAL4*-induced *Ras^V12^, scrib^RNAi^*tumour*, QF2* induced *Ras^V12^, scrib^RNAi^* tumours were able to induce a 40% reduction in muscle/cuticle attachment (Dark et al., 2022; Lodge et al., 2021) (Figure 2 A-E). Upon overexpression of Akt in the fatbody (*r4-GAL4*) of tumour bearing animals (*QRas^V12^, scrib^RNAi^*), we found there was a significant improvement in muscle morphology (Figure 2 F-H). Similarly, overexpression of *InR^CA^*in the fatbody of tumour bearing animals (*QRas^V12^, scrib^RNAi^*) also improved muscle integrity (Figure 2 I-K). Together, our data indicate that the elevation of fatbody insulin signalling level can improve muscle integrity in the context of cachexia.

**Figure 2:**
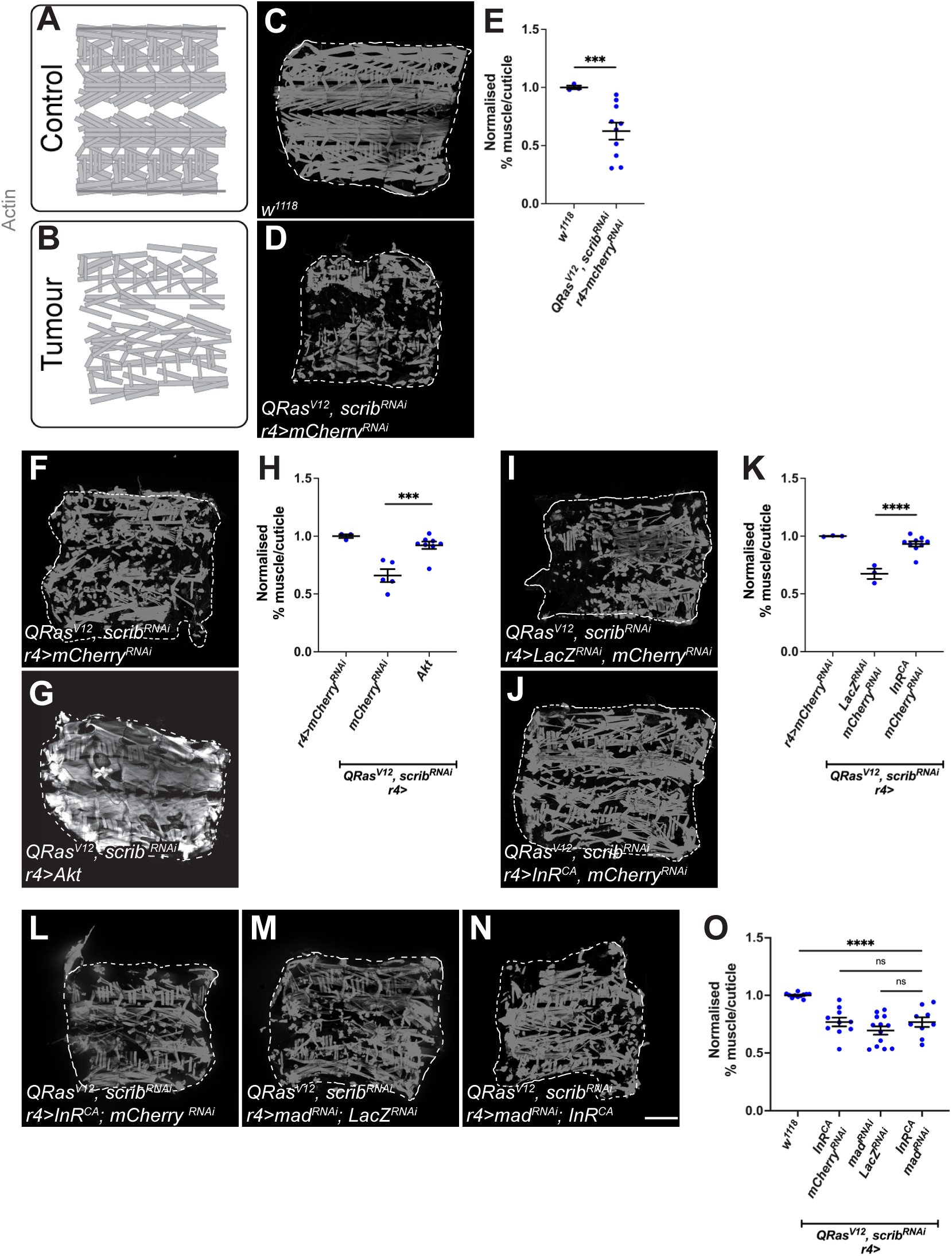
Insulin signalling in the fatbody improves muscle integrity in cachectic animals. (A-B) Schematic depicting intact muscle fillet in control animals, versus deteriorated muscle fillets in tumour animals. (C-D) Muscle fillets stained with phalloidin (Actin) from *w^1118^* or *QRas^V12^scrib^RNAi^* tumour-bearing animals where *mCherry^RNAi^* was expressed in the fatbody (*r4-GAL4*). Detachments are marked with yellow arrows. (E) Quantification of normalized muscle detachment in C-D. *w^1118^* (n=3), *QRas^V12^scrib^RNAi^* (n=10). (F-G) Muscle fillets stained with phalloidin (Actin) from *QRas^V12^scrib^RNAi^*tumour-bearing animals raised at 25°C, where in the fatbody, either *mCherry^RNAi^*, *or Akt was* overexpressed. (H) Quantification of normalized muscle detachment in F-G, *r4>mCherry^RNAi^ (n=3)*. *QRas^V12^scrib^RNAi^; r4> mCherry^RNAi^* (n=5), *QRas^V12^scrib^RNAi^; r4>Akt* (n=8). (I-J) Muscle fillets stained with phalloidin (Actin) from *QRas^V12^scrib^RNAi^* tumour-bearing animals raised at 25°C, where in the fatbody, either *lacZ;mCherry^RNAi^*, *or InR^CA^; mCherry^RNAi^*was overexpressed. (K) Quantification of normalized muscle detachment in I-J, *r4>mCherry^RNAi^ (n=3)*. *QRas^V12^scrib^RNAi^; r4>lacZ;mCherry^RNAi^* (n=3), *QRas^V12^scrib^RNAi^; r4>InR^CA^; mCherry^RNAi^* (n=9). (L-N) Muscle fillets of *QRas^V12^scrib^RNAi^*tumour-bearing animals raised at 25°C where *InR^CA^*; *mCherry^RNAi^* or *mad^RNAi^; LacZ^RNAi^* or *InR^CA^; mad^RNAi^* was expressed in the fatbody (*r4-GAL4*). (O) Quantification of muscle detachment in P-R. *w^1118^* (n=3), *InR^CA^*; *mCherry^RNAi^* (n=8) or *mad^RNAi^; LacZ^RNAi^* (n=14) or *InR^CA^; mad^RNAi^* (n=9). Scale bar is 50µm for fatbody pAkt and pMad staining Graphs are represented as Mean ± SEM, n= the number of samples. *(*) p<0.05 (**) p< 0.01, (***) p< 0.001, (****) p < 0.0001, (ns) p>0.05*

To test if fatbody insulin signalling facilitates cachexia progression via TGF-β signalling, we next activated insulin signalling via fatbody expression of *InR^CA^* and simultaneously knocked down TGF-β signalling via fatbody expression of *mad^RNAi^* (Figure 2 L-O) in tumour bearing animals (*QRas^V12^, scrib^RNAi^*). We have previously shown that *mad^RNAi^* was sufficient to rescue muscle integrity (Lodge et al., 2021), and here found that *InR^CA^* (*InR^CA^;mCherry^RNAi^)* was similarly able to rescue muscle integrity (Figure 2 I-K). Co-expression of *InR^CA^* and *mad^RNAi^* gave a similar rescue as either *InR^CA^* (*InR^CA^;mCherry^RNAi^*) or *mad^RNAi^ (mad^RNAi^;LacZ^RNAi^)* alone, suggesting that TGF-β signalling likely acts downstream of insulin signalling in the fatbody of tumour-bearing animals.

Fatbody insulin signalling has been known to modulate the overall size of the animal by altering systemic ecdysone levels and thus we considered whether the insulin signalling pathway may influence cachexia via this growth hormone (Caldwell et al., 2005; Colombani et al., 2005; Mirth et al., 2005). We have previously shown that decreasing systemic ecdysone levels, via prothoracic gland (PG)-specific expression of an RNAi against *torso*, increased overall body size, but was not sufficient to cause muscle detachment (Lodge et al., 2021). Here we show that a reduction in body size by increasing global ecdysone levels through PG-specific expression of *Ras^V12^* (Caldwell et al., 2005) failed to alter the level of muscle detachment in tumour bearing animals (Figure S1 A-C, *QRas^V12^, scrib^RNAi^*). Finally, in tumour bearing animals fed a sterol-free diet, that underwent a prolonged 3^rd^ instar stage due to reduced ecdysone levels (Parkin and Burnet, 1986), we activated insulin signalling in the fatbody via Akt overexpression (*QRas^V12^, scrib^RNAi^*). We found that this manipulation caused a significant decrease in pMad levels in the fatbody and a rescue of muscle detachment (Figure S1 D-I), similar to animals fed a standard diet (Figure 1 O-Q, Figure 2 F-H). Together, our data suggest that systemic ecdysone levels are unlikely to be involved in modulating tumour-induced muscle detachment or to mediate the role of fatbody Insulin signalling in regulating muscle detachment.

### Tumour secreted ImpL2 and Gbb act additively to affect muscle integrity

Although our data suggest that insulin and TGF-β signalling act in a linear pathway in the fatbody to facilitate cachexia, it is unclear how the tumour-secreted proteins ImpL2 and Gbb interact with each other to facilitate the progression of cachexia. To determine if the two proteins interact, we first assessed whether ImpL2 regulates TGF-β signalling by influencing *gbb* expression in the tumour. Upon the knockdown of *Impl2*, we found that tumour *gbb* was not significantly altered (Figure S3A). Next, to determine whether ImpL2 and Gbb acted synergistically to affect muscle integrity, we knocked down both *gbb* and *ImpL2* in the tumour (*Ras^V12^, dlg^RNAi^*). Inhibition of both ligands resulted in a greater rescue in muscle integrity than knockdown of either ligand alone (Figure S2 A-E). We suspected that these additive effects of the proteins were because each protein rescued different aspects of cachexia, and thus tested their effects on two important aspects of cachexia: protein synthesis and ECM accumulation. Protein translation (measured using the O-propargyl-puromycin incorporation assay, OPP assay) is significantly downregulated during cachexia, and ECM proteins such as Nidogen accumulated in the fatbody of tumour bearing animals (Lodge et al., 2021, Figure S2 F-H, N-P). We next examined whether the knockdown of *gbb* and *ImpL2*, either alone or together in the tumour, rescued protein synthesis or ECM accumulation. Knockdown of *gbb* alone in the tumour did not significantly rescue protein synthesis in the fatbody, however, *ImpL2^RNAi^* alone or *ImpL2^RNAi^* and *gbb^RNAi^* together rescued protein synthesis (Figure S2 I-M). Conversely, knockdown of *gbb* alone or knockdown of *gbb* together with *ImpL2* significantly rescued the Nidogen overaccumulation defects observed at the plasma membrane of fatbody from tumour-bearing animals, while *ImpL2^RNAi^* alone did not (Figure S2 Q-U). Finally, co-knockdown of *gbb* and *ImpL2* in the tumour significantly rescued the reduction in OPP and Nidogen levels observed in the muscles of tumour-bearing animals (Figure S3 B-I). Altogether, our data indicate that *ImpL2^RNAi^* and *gbb^RNAi^* rescue different aspects of cachexia to additively rescue muscle degradation.

### Muscle insulin signalling regulates atrophy and muscle integrity in cachexia

A key feature of cancer-induced wasting is the reduction in myofiber size, termed atrophy. This reduction in muscle size can be measured as a ratio of muscle width/length of the VL 3 muscle (Graca et al., 2022; Lodge et al., 2021). Since tumour specific *ImpL2* inhibition significantly rescued muscle translation (Figure S2-S3), we suspected that insulin signalling in the muscle may play a role in modulating muscle integrity. To test this, we activated insulin signalling in the muscle of tumour bearing animals (*QRas^V12^, scrib^RNAi^*) by overexpressing *Akt* with the muscle specific driver *MHC-GAL4*. This manipulation significantly rescued muscle integrity (Figure S4 A-C) and muscle atrophy (Figure S4 D-F), without affecting muscle ECM levels (Figure S4 G-H). This suggests that muscle insulin signalling predominantly regulate translation and atrophy. Interestingly, while muscle pMad levels are elevated in tumour-bearing animals and that tumour-specific *ImpL2* inhibition was able to reduce muscle TGF-β signalling in tumour-bearing animals (Figure S4 P-R); muscle specific expression of *mad^RNAi^*in tumour bearing animals (*QRas^V12^, scrib^RNAi^*) using *MHC-GAL4* (Figure S4 S) or *Mef2-GAL4* (data not shown) was not able to improve muscle integrity. This suggests that although muscle TGF-β signalling is responsive to circulating insulin levels, muscle TGF-β signalling is functionally dispensable in facilitating muscle degradation during cachexia.

### Fatbody TGF-β signalling, Rab10 and SPARC regulate ECM accumulation and muscle integrity in cachexia

Next, we examined whether modulating fatbody insulin or TGF-β signalling can improve muscle integrity. We found that fatbody specific Akt expression was not able to rescue muscle atrophy caused by *QRas^V12^Scrib^RNA^*^i^ tumours (Figure S4 J-L). As TGF-β signalling acts downstream of insulin signalling in the fatbody of tumour bearing animals, we next asked if inhibition of TGF-β signalling in the fatbody via the expression of an RNAi against *mad* improved muscle atrophy in tumour bearing animals (*QRas^V12^Scrib^RNA^*^i^). Interestingly, although fatbody *mad^RNAi^* expression was able to improve overall muscle integrity (Lodge et al., 2021), this manipulation did not significantly improve muscle atrophy (Figure S4 M-O). It however was able to significantly reduced ECM accumulation in the fatbody (Figure 3 A-B, E) which consequently also increased muscle Nidogen levels (Figure 3 C-D, F). Together, this data suggests that fatbody insulin and TGF-β signalling converge to specify ECM levels in the fatbody and in turn, the muscle during cachexia.

**Figure 3:**
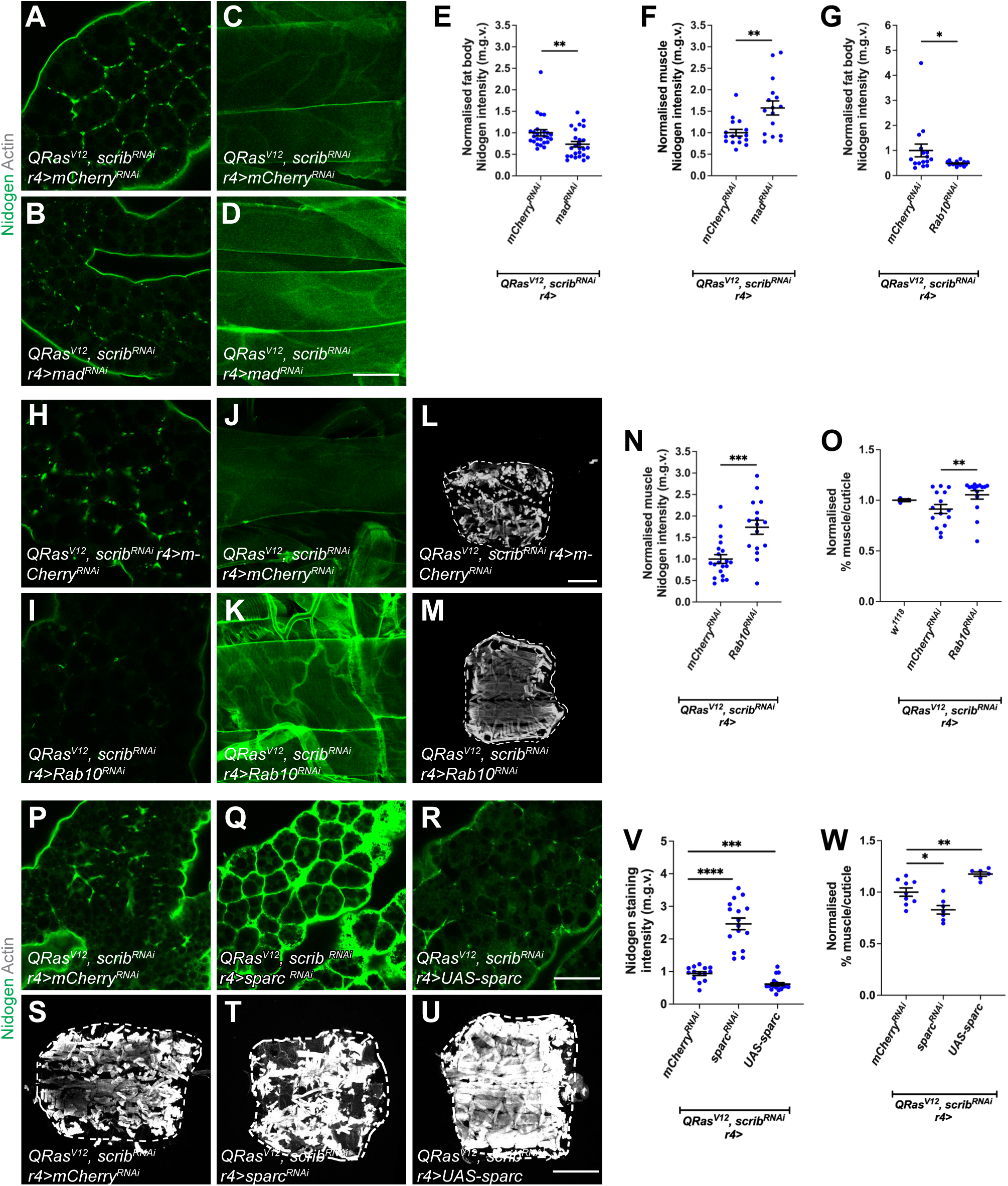
TGF-ß signaling in the fatbody rescues muscle detachment via the regulation of fatbody ECM secretion. (A-B) Fatbody stained with the ECM protein nidogen from *QRas^V12^scrib^RNAi^* tumour-bearing animals, where *mCherry^RNAi^*or *mad^RNAi^* was expressed in the fatbody (*r4-GAL4*). (E) Quantification of fatbody nidogen staining in A-B. *mCherry^RNAi^* (n=25), *mad^RNAi^* (n=25). (C-D) Muscles stained with nidogen from *QRas^V12^scrib^RNAi^*tumour-bearing animals, where *mCherry^RNAi^* or *mad^RNAi^* was expressed in the fatbody (*r4-GAL4*). (F) Quantification of muscle nidogen in C-D. *mCherry ^RNAi^* (n=16), *mad^RNAi^* (n=16). (H-I) Fatbody stained with the ECM protein nidogen from *QRas^V12^scrib^RNAi^* tumour-bearing animals, where *mCherry^RNAi^*or *Rab10^RNAi^* was expressed in the fatbody (*r4-GAL4*). (G) Quantification of normalized fatbody nidogen staining in H-I. *mCherry^RNAi^* (n=16), *Rab10^RNAi^* (n=16). (J-K) Muscles stained with nidogen from *QRas^V12^scrib^RNAi^* tumour-bearing animals, where *mCherry^RNAi^* or *Rab10^RNAi^* was expressed in the fatbody (*r4-GAL4*). (N) Quantification of normalized muscle nidogen staining in J-K. *mCherry^RNAi^* (n=20), *Rab10^RNAi^* (n=16). (L-M) Muscle fillets stained with phalloidin (Actin) from *QRas^V12^scrib^RNAi^* tumour-bearing animals, where *mCherry^RNAi^*or *Rab10^RNAi^* was expressed in the fatbody (*r4-GAL4*). Day 6 muscles used here. (O) Quantification of normalized muscle detachment in L-M. *w^1118^* (n=3), *QRas^V12^scrib^RNAi^; r4> mCherry^RNAi^* (n=15), *QRas^V12^scrib^RNAi^;r4> Rab10^RNAi^* (n=15). (P-R) Fatbody stained with the ECM protein nidogen from *QRas^V12^scrib^RNAi^* tumour-bearing animals, where *mCherry^RNAi^*or *sparc^RNAi^* or *UAS-sparc* was expressed in the fatbody (*r4-GAL4*). (V) Quantification of normalized fatbody nidogen staining in P-R. *mCherry^RNAi^* (n=12), *sparc^RNAi^* (n=16), *UAS-sparc* (n=10). (S-U) Muscle fillets stained with phalloidin (Actin) from *QRas^V12^scrib^RNAi^* tumour-bearing animals, where *mCherry^RNAi^*or *sparc^RNAi^* or *UAS-sparc* was expressed in the fatbody (*r4-GAL4*). (V) Quantification of normalized muscle fillet in S-U. *QRas^V12^scrib^RNAi^; r4> mCherry^RNAi^* (n=10), *QRas^V12^scrib^RNAi^; r4>sparc^RNAi^* (n=6), *QRas^V12^scrib^RNAi^; r4>UAS-sparc* (n=6).

It has previously been shown that muscle ECM proteins are mostly derived from the fatbody and blood cells (Dai et al., 2018), furthermore, in the context of cachexia, we observed a correlation between ECM accumulation in the fatbody, and ECM depletion in the muscle. We therefore hypothesised that cachectic fatbody may be trapping ECM proteins and preventing ECM secretion to the muscle, causing muscle degradation, To explore whether this may be the case, we blocked endocytosis in the fatbody (*CG-GAL4*) through expression of a temperature sensitive dominant negative form of *shibire* (*shi^t^*^s^) (Zang et al., 2015). It was previously shown that fatbody *shibire* knockdown causes trapping of outgoing ECM proteins, such as Nidogen, in the fatbody membrane (Zang et al., 2015). Consistent with our hypothesis, we found that blocking endocytosis resulted in a significant downregulation of Nidogen in the muscles (Figure S5 A-C) and an increase in muscle detachment (Figure S5 D-F). Together, these data suggest that fatbody ECM accumulation may contribute to the muscle ECM deficit and muscle detachment.

To test whether ECM accumulation in the fatbody can affect muscle detachment in the context of cancer cachexia, we next tried to modulate ECM levels in the fatbody in tumour bearing animals (*QRas^V12^Scrib^RNA^*^i^). It was previously reported that the overexpression of a small GTPase protein called Rab10 can cause an accumulation of ECM proteins (Isabella and Horne-Badovinac, 2016). To specifically prevent the accumulation of ECM proteins, we overexpressed *Rab10^RNAi^* in the fatbody of tumour bearing animals. This manipulation significantly reduced the accumulation of fatbody ECM protein Nidogen (Figure 3 G-I), increased muscle Nidogen levels (Figure 3 J-K, N), and improved muscle attachment (Figure 3 L-M, O), suggesting that modulating fatbody ECM levels in cachectic animals can have a direct effect on muscle ECM levels and integrity.

We next investigated whether modulating SPARC, the *Drosophila* homolog of BM40/SPARC/osteonectin, that is known to be required for ColIV secretion, can affect muscle integrity in cancer cachexia. It has been shown previously that the loss of SPARC caused the retention of ColIV in the membranes of fatbody cells (Pastor-Pareja and Xu, 2011; Shahab et al., 2015). Consistent with this, the expression of a *sparc* RNAi in the fatbody of tumour bearing animals caused a significant accumulation of Nidogen, and further deterioration of muscle morphology (Figure 3 Q, T, V-W). Conversely, overexpression of *sparc* (Portela et al., 2010), significantly reduced Nidogen accumulation in the fatbody, and significantly improved muscle morphology (Figure 3 R, U, V-W). Finally, we asked whether increasing Collagen IV (*viking* (*vkg*) and *Collagen at 25C* (*Cg25C*)) (Pastor-Pareja and Xu, 2011) levels in the fatbody of tumour bearing animals can improve muscle integrity (Figure S5 G-I). Interestingly, this manipulation was also able to improve muscle morphology. Together, our data suggest that fatbody ECM accumulation contributes to a muscle ECM deficit, as well as muscle detachment in cancer cachexia.

### Insulin signalling can function via TGF-β signalling in the wildtype fatbody

Next, we assessed whether the cross-regulatory relationship between insulin signalling and TGF-β signalling observed in the cachectic fatbody is relevant in a developmental context. For this, we first starved larvae by subjecting animals to a diet of 1% agarose in PBS for 24hrs to reduce their levels of circulating Ilps (Brogiolo et al., 2001). Starved larvae exhibited the expected downregulation in fatbody pAkt (data not shown) and a significant increase in fatbody pMad levels (Figure 4A-C) compared to controls fed on a standard laboratory diet. Next, we used *CG-GAL4* to express a constitutive activate form of *InR* (*InR^CA^*). This caused an upregulation in fatbody pAkt (data not shown) and significant reduction in fatbody pMad levels (Figure 4D-F). Conversely, when *CG-GAL4* was used to overexpress the gene for the adaptor protein *p60*, which acts downstream of InR to recruit DP110 and downregulates insulin signalling when overexpressed (Weinkove et al., 1999), we observed an upregulation in fatbody pMad levels (Figure 4 G-I). Together, our data suggest that the insulin signalling pathway inhibits TGF-β signalling in the wildtype fatbody.

**Figure 4:**
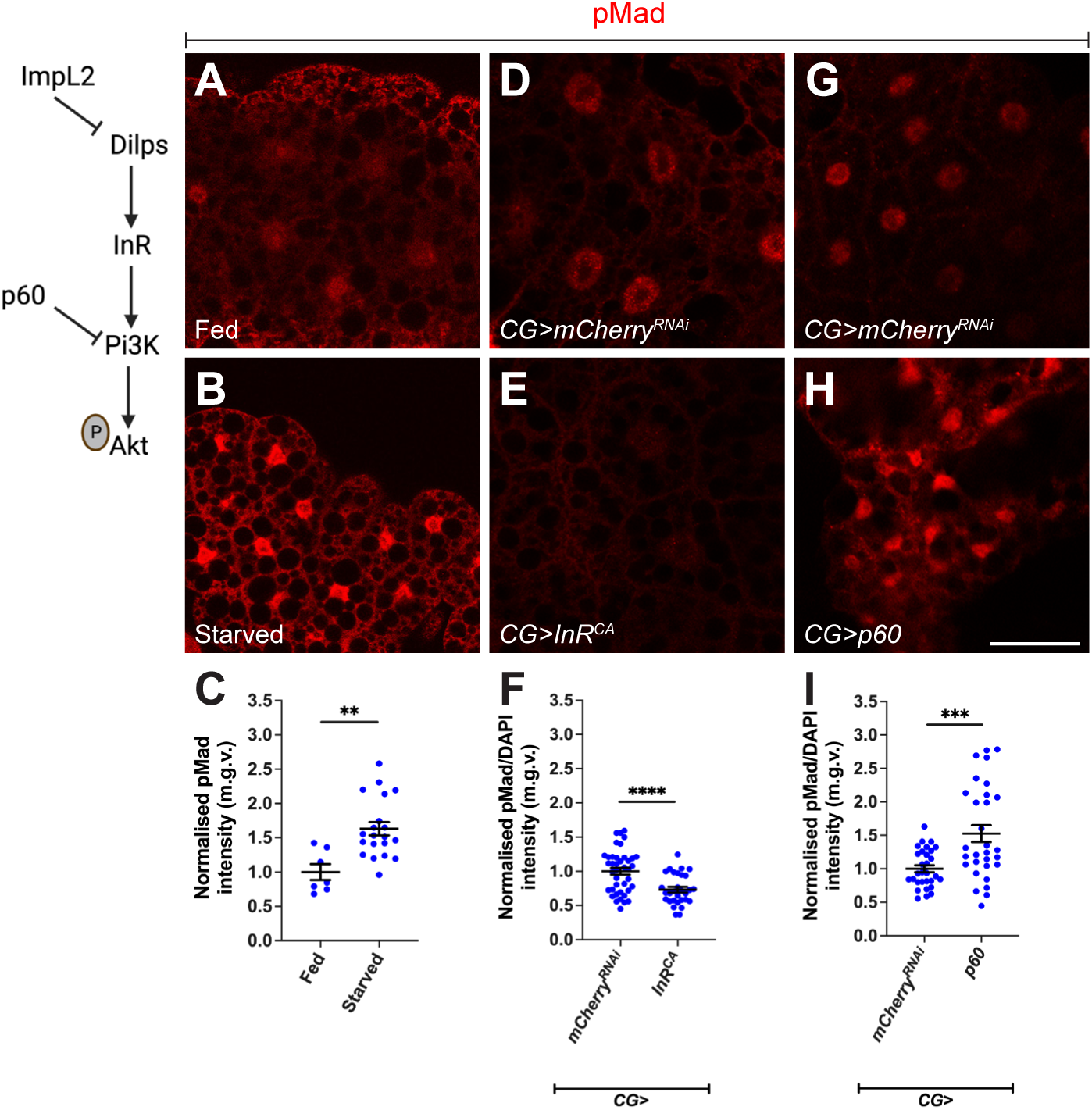
Fatbody insulin signaling negatively regulates TGF-ß signaling. (A-B) Fatbodies from fed and starved animals with TGF-ß signaling activation indicated by pMad staining. (C) Quantification of normalized pMad staining in A-B. Fed (n=7), starved (n=20). (D-E) Fatbody from animals raised at 18°C and expressing *mCherry^RNAi^* or *InR^CA^* under the control of *CG-GAL4*, with TGF-ß signaling activation indicated by pMad staining. (F) Quantification of normalized pMad staining in D-E. *mCherry^RNAi^*(n=30), *InR^CA^* (n=40). (G-H) Fatbody from animals expressing *mCherry^RNAi^* or *p60* under the control of *CG-GAL4*, with TGF-ß signaling activation indicated by pMad staining. (I) Quantification of normalized pMad staining in G-H. *mCherry^RNAi^* (n=30), *p60* (n=30). Scale bar is 50µm. Graphs are represented as Mean ± SEM, n= the number of samples. *(*) p<0.05 (**) p< 0.01, (***) p< 0.001, (****) p < 0.0001, (ns) p>0.05*

We next tested if reduced Insulin signalling in the fatbody phenocopies the Vkg accumulation observed upon fatbody activation of TGF-β signalling. Vkg accumulated in the membranes of fatbody where *p60* was overexpressed using *r4-GAL4* (Figure 5 A-C). Similarly, Vkg accumulation was observed upon the overexpression of a dominant negative form of *Target of Rapamycin* (*TOR^DN^*) using *r4-GAL4*, which is activated downstream of pAkt in the insulin signalling cascade (Figure 5 E-G). Moreover, as with the activation of Tkv, expression of *TOR^DN^* did not induce muscle detachment (Figure 5 D). This indicates that these signals are indeed functionally equivalent and thus wild-type fatbody insulin signalling may also act through TGF-β. Interestingly, however, fatbody *p60* overexpression caused a small but significant reduction in muscle attachment, suggesting that fatbody insulin signalling may play additional roles to induce muscle detachment that are independent of TGF-β (Figure 5 H).

**Figure 5:**
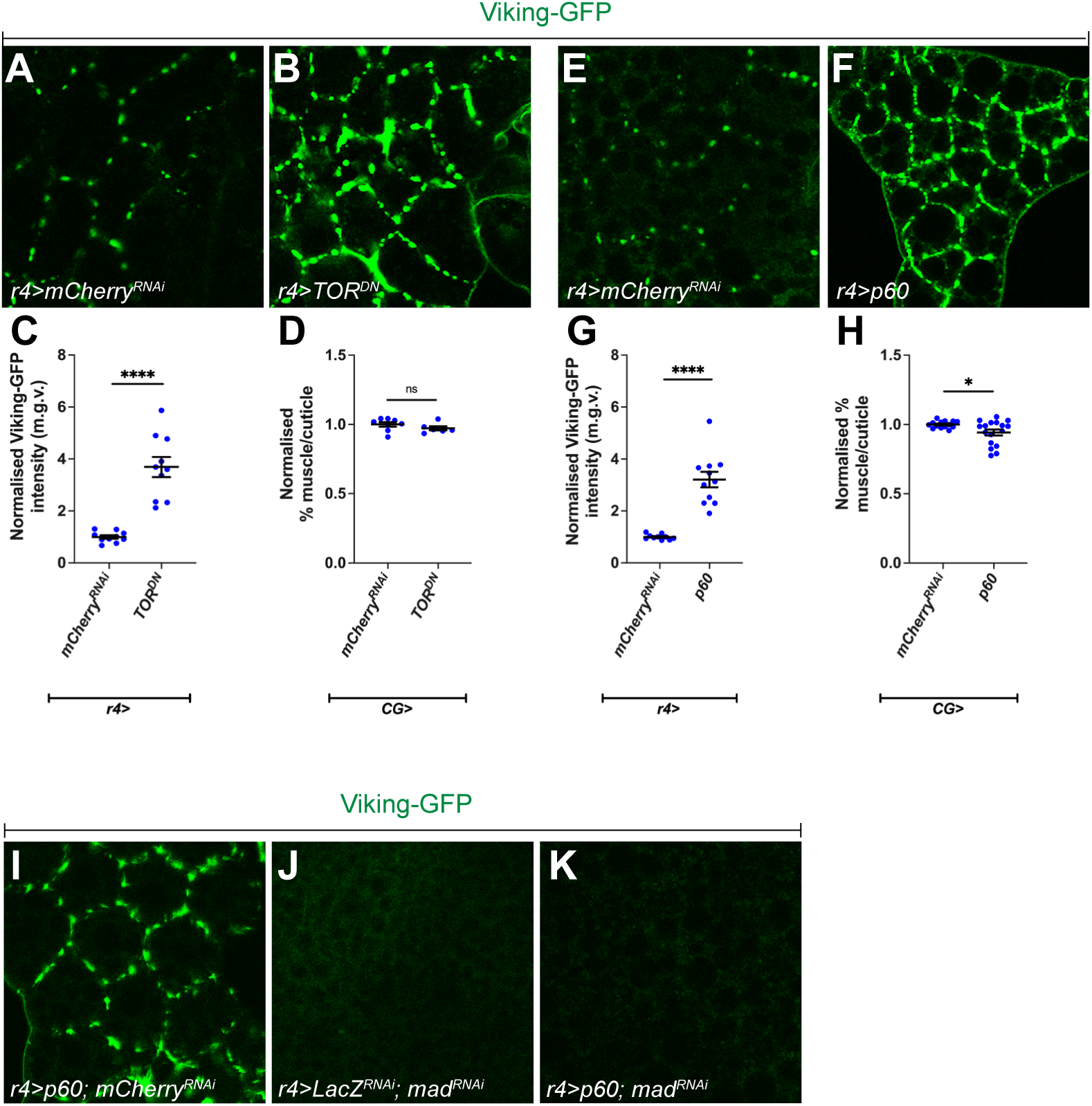
Fatbody insulin signaling negatively regulates ECM accumulation. (A-B) Viking-GFP localization in fatbodies from animals expressing *mCherry^RNAi^* or *TOR^DN^* under the control of *r4-GAL4*. (C) Quantification of normalized Viking-GFP intensity in A-B*. mCherry^RNAi^* (n=10) *TOR^DN^* (n=9). (D) Quantification of normalized muscle detachment caused by the expression of *mCherry^RNAi^* or *TOR^DN^* under the control of *r4-GAL4. mCherry^RNAi^* (n=8) *TOR^DN^* (n=7). (E-F) Viking-GFP localization in fatbody from animals expressing *mCherry^RNAi^* or *p60* under the control of *r4-GAL4*. (G) Quantification of normalized Viking-GFP intensity in E-F. *mCherry^RNAi^* (n=9), *p60* (n=10). (H) Quantification of normalized muscle detachment caused by the expression of *mCherry^RNAi^* or *p60* under the control of *r4-GAL4. mCherry^RNAi^* (n=12), *p60* (n=16). (I-K) Viking-GFP localization in fatbody from animals expressing *p60; mCherry^RNAi^* or *lacZ^RNAi^; mad^RNAi^* or *p60; mad^RNAi^* under the control of *r4-GAL4*. Scale bar is 50µm for fatbody staining and 50µm for muscle Viking staining. Graphs are represented as Mean ± SEM, n= the number of samples. *(*) p<0.05 (**) p< 0.01, (***) p< 0.001, (****) p < 0.0001, (ns) p>0.05*

To determine whether insulin signalling knockdown induces ECM accumulation in the fatbody via TGF-β signalling, we expressed *p60* while simultaneously knocking down TGF-β signalling using *mad^RNAi^*. Similar to *CG>lacZ^RNAi^; mad^RNAi^*, very little Vkg was detected at the plasma membrane of *CG>p60*; *mad^RNAi^* fatbodies (Figure 5 I-K). Together, these data suggest that the accumulation of ECM proteins in the fatbodies of *CG>p60* or *CG>TOR^DN^* larvae is dependent on *mad*, functionally placing insulin signalling upstream of TGF-β signalling in the wild-type fatbody (Figure 5 I-K).

### Insulin signalling affects TGF-β signalling via TOR and S6K

pAkt triggers phosphorylation cascades that result in deactivation of the transcription factor Forkhead box, sub-group O (FOXO) and activation of TOR. Once activated, TOR activates S6K and deactivates 4E-BP respectively by inducing their phosphorylation(Zhang et al., 2000). Together, FOXO, TOR, S6K and 4E-BP make up the most downstream components of the insulin signalling pathway and induce the cellular changes triggered by the activation of the pathway (Giannakou and Partridge, 2007; Semaniuk et al., 2021). To test which of these factors regulate TGF-β signalling, we used *CG-GAL4* to overexpress *FOXO*, *TOR^DN^*, or constitutively activated forms of *S6K* (*S6K^CA^*) or *4E-BP* (*4E-BP^CA^*) in the fatbody. We found that fatbody from *CG*>*4E-BP^CA^*and *CG>FOXO* larvae exhibited no change in pMad levels (Figure 6 D-F, J-L), whereas fatbody from *CG>TOR^DN^* larvae exhibited increased pMad levels (Figure 6A-C) and *CG*>*S6K^CA^* fatbody exhibited reduced pMad levels (Figure 6 G-I). To assess whether *InR* inhibited TGF-β signalling in the fatbody via *TOR*, we tested the epistatic relationship between *TOR^DN^* and *InR^CA^* expression. We found that the co-expression of *TOR^DN^*and *InR^CA^* caused elevated pMad, phenocopying knockdown of *TOR* (*TOR^DN^*) alone, suggesting that insulin signalling modulates fatbody TGF-β signalling via *TOR* (Figure 6M-Q).

**Figure 6:**
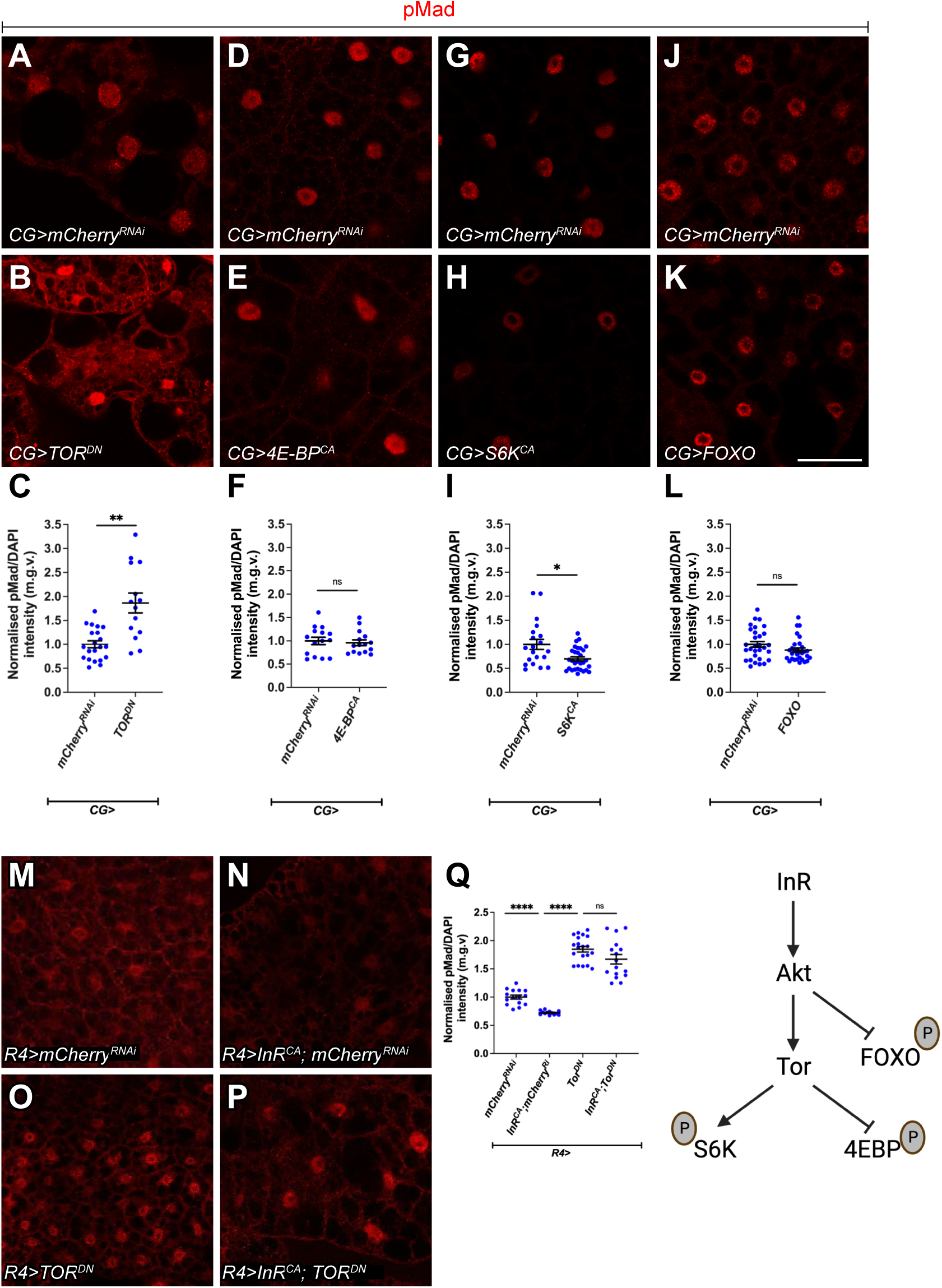
Fatbody TOR signaling negatively regulates TGF-ß signaling. (A-B) Fatbody from animals expressing *mCherry^RNAi^* or *TOR^DN^* under the control of *CG-GAL4*, with TGF-ß signaling activation indicated by pMad staining. (C) Quantification of normalized pMad staining in A-B. *mCherry^RNAi^* (n=21), *TOR^DN^* (n=14). (D-E) Fatbodies from animals expressing *mCherry^RNAi^* or *4E-BP^CA^*under the control of *CG-GAL4*, with TGF-ß signaling activation indicated by pMad staining. (F) Quantification of normalized pMad staining in D-E. *mCherry^RNAi^* (n=15), *4E-BP^CA^* (n=15). (G-H) Fatbody from animals expressing *mCherry^RNAi^* or *S6K^CA^* under the control of *CG-GAL4*, with TGF-ß signaling activation indicated by pMad staining. (K) Quantification of normalized pMad staining in G-H. *mCherry^RNAi^* (n=20), *S6K^CA^* (n=30). (J-K) Fatbody from animals expressing *mCherry^RNAi^* or *FOXO* under the control of *CG-GAL4*, with TGF-ß signaling activation indicated by pMad staining. (L) Quantification of normalized pMad staining in J-K. *mCherry^RNAi^* (n=30), *FOXO* (n=30). (M-P) Fatbody from animals expressing *mCherry^RNAi^* or *Tor^DN^ or InR^CA^*; *mCherry^RNAi^* or *InR^CA^;Tor^DN^*under the control of *r4-GAL4*, with TGF-ß signaling activation indicated by pMad staining. (Q) Quantification of normalized pMad staining in M-P. *mCherry^RNAi^* (n=16), *Tor^DN^* (n=20), *InR^CA^*; *mCherry^RNAi^* (n=12), *InR^CA^;Tor^DN^*(n=16). The same *mCherry^RNAi^* and *InR^CA^*; *mCherry^RNAi^* data points were used as in Figure 7F. Scale bar is 50µm. Graphs are represented as Mean ± SEM, n= the number of samples. *(*) p<0.05 (**) p< 0.01, (***) p< 0.001, (****) p < 0.0001, (ns) p>0.05*

### Insulin does not regulate TGF-β signalling in the muscle or the wing imaginal discs

To determine whether the effect of insulin signalling on TGF-β signalling is fatbody specific, we next assessed whether modulating insulin signalling in the muscle, or the wing discs affects pMad levels in these tissues. We found that overexpression of a constitutively active form of *DP110* (*DP110^CAAX^*) (Leevers et al., 1996) using the muscle driver *MHC-GAL4* did not significantly alter muscle pMad staining (Figure S6 A-C). In the wing imaginal disc, heat shock-induced clones overexpressing *InR^CA^*or *p60* exhibited expected changes in clone size (Figure S6 D-E’) and pAkt levels (data not shown). However, these alterations in insulin signalling did not significantly alter TGF-β signalling (Figure S6 D-E’). Similarly, *tkv^CA^* or *mad^RNAi^* clones caused altered clone size (Figure S6 F-G’) and changes in pMad levels (data not shown) but caused no significant changes in pAkt levels in the wing (Figure S6 F-G’). This indicates that the cross-regulation of these signalling pathways does not hold in the muscle or the wing imaginal discs.

### Insulin signalling influences TGF-β signalling by modulating fatbody *sog* expression

Since the Bone Morphogenic Protein (BMP) arm of the TGF-β signalling pathway controls Mad phosphorylation, we suspected that the insulin signalling pathway may modulate TGF-β signalling levels by directly modulating the transcription of components or regulators of the BMP signalling pathway. To test this, qPCR was conducted on the fatbody of *CG>InR^CA^* and *CG>mCherry^RNAi^* animals. We assessed the expression levels of the BMP ligand encoding genes (*gbb*, *decapentaplegic* [*dpp*] and *maverick* [*mav*], but not *scarecrow* as it is only expressed in the CNS during the larval stage (Yoo et al., 2020)), the four BMP receptor encoding genes (*thickveins* [*tkv*], *saxophone* [*sax*], *punt* and *wishful thinking* [*wit*]), the BMP signalling inhibitor *sog*, the transcription factor *mad*, and the putative regulator of Mad dephosphorylation *dullard* (Urrutia et al., 2016). Since non-BMP components of the TGF-β signalling pathway have also been observed to influence Mad phosphorylation (Gesualdi and Haerry, 2007; Peterson et al., 2012), we also assessed the transcription of several other TGF-β signalling pathway genes: *babo*, *myoglianin* (*myo*), *dawdle* (*daw*) and *smad on X* (*smox*). Strikingly, we found that *sog* transcription was increased 2-fold in the fatbody of *CG*>*InR^CA^*larvae, while none of the other genes tested exhibited any changes in transcription levels (Figure 7A). To assess whether Sog mediates the downstream effects of insulin signalling on TGF-β, we expressed *sog^RNAi^* in the fatbody of larvae expressing *InR^CA^* and tested whether *sog* knockdown inhibits the effects of *InR^CA^* on fatbody pMad levels (Figure 7B-F). We found that *sog* knockdown increased pMad level, and the expression of *sog^RNAi^* together with *InR^CA^*resulted in an increase in pMad level similar to *sog^RNAi^* overexpression alone. These findings suggest that Sog lies downstream of insulin signalling and mediates the effects of insulin signalling on TGF-β signalling.

**Figure 7:**
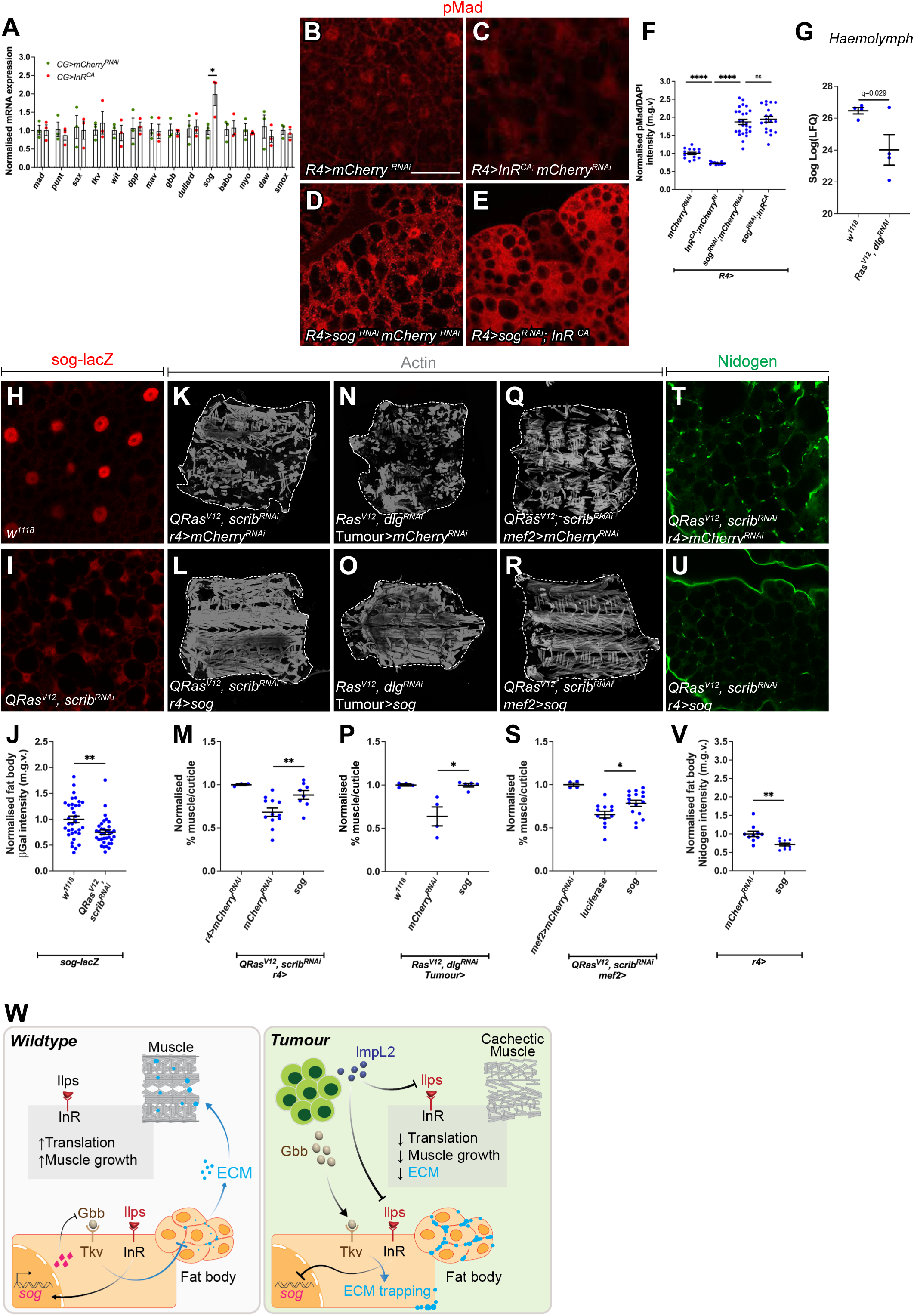
Insulin signaling activates the TGF-ß inhibitor Sog which in turn regulates muscle detachment during cancer cachexia. (A) Fatbody qPCRs showing mRNA expression levels of TGF-ß receptors and ligands in *InR^CA^* or *mCherry^RNAi^* larvae (raised at 18°C) with *CG-GAL4* (n=3). (B-E) Overexpression of *mCherry^RNAi^, InR^CA^; mCherry^RNAi^, Sog^RNAi^; mCherry^RNAi^ and Sog^RNAi^;InR^CA^* under the control of *r4-GAL4* stained for pMad. (F) Quantification of pMad levels of B-E. *mCherry^RNAi^* (n=17), *InR^CA^; mCherry^RNAi^*(n=7)*, Sog^RNAi^; mCherry^RNAi^* (n=26), *and Sog^RNAi^;InR^CA^*(n=19), The same *mCherry^RNAi^* and *InR^CA^*; *mCherry^RNAi^* data points were used as in Figure 6Q. (G) Sog levels in the haemolymph of *w^1118^* or *Ras^V12^Dlg^RNAi^* tumour-bearing animals (n=4). (H-I) sog-lacZ staining in *w^1118^*or *QRas^V12^scrib^RNAi^* fatbody. (J) Quantification of sog-lacZ levels in H-I. *w^1118^*(n=36), *QRas^V12^scrib^RNAi^*(n=39). (K-L) Muscle fillets stained with phalloidin (Actin) from animals where *mCherry^RNAi^* or *sog* was expressed in the fatbody using *r4-GAL4*. (M) Quantification of muscle detachment in K-L. *w^1118^* (n=3), *mCherry^RNAi^* (n=10), *sog* (n=8). (N-O) Muscle fillets stained with phalloidin (Actin) from upon tumour-specific overexpression of *mCherry^RNAi^* or *Sog* in *Ras^V12^Dlg^RNAi^* tumour-bearing animals. (P) Quantification of muscle detachment in N-O. *w^1118^* (n=3), *mCherry^RNAi^* (n=4), *sog* (n=6). (Q-R) Muscle fillets stained with phalloidin (Actin) from upon tumour-specific overexpression of *luciferase* or *sog* in the muscle via *MHC-GAL4* (S) Quantification of muscle detachment in Q-R. *w^1118^* (n=3), *luciferase* (n=10), *Sog* (n=16). Scale bar is 500µm. (T-U) Fatbody stained with the ECM protein nidogen from *QRas^V12^scrib^RNAi^* tumour-bearing animals, where *mCherry^RNAi^*or *UAS-sog* was expressed in the fatbody (*r4-GAL4*). (V) Quantification of normalized fatbody nidogen staining in T-U. *mCherry^RNAi^* (n=9), *UAS-sog* (n=10). Scale bar is 50µm for fatbody. (W) Summary: <u>Left</u>: During development, insulin signaling in the fatbody activates the transcription of *sog*, which inhibits Gbb and prevents the activation of TGF-ß signaling in the fatbody. This allows fatbody ECM to be secreted to function in the muscle. Insulin signaling in the muscle in parallel enhances translation and muscle growth. <u>Right:</u> In Tumour bearing/cachectic animals, Tumours secrete two ligands: ImpL2 and Gbb. In the fatbody, ImpL2 inhibits insulin signalling, preventing the transcription of *sog* and thus Sog can no longer inhibit Gbb. In addition, tumour secreted Gbb binds to Tkv to activate TGF-ß signaling in the fatbody, resulting in an accumulation of ECM proteins, to prevent ECM transport out of the fatbody to reach the muscle. In the muscle, ImpL2 inhibits insulin signalling, which inhibits translation and muscle growth. Graphs are represented as Mean ± SEM, n= the number of samples. *(*) p<0.05 (**) p< 0.01, (***) p< 0.001, (****) p < 0.0001, (ns) p>0.05*

### Sog is a critical regulator of cachexia

Since insulin signalling likely regulates TGF-β signalling by modulating *sog* expression, we assessed whether Sog modulation is relevant in cachectic animals. Analysis of our previously published hemolymph proteomics data (Lodge et al., 2021) showed that Sog is one of the most downregulated circulating proteins in tumour-bearing animals (Figure 7G). Using a lacZ enhancer trap line of *sog*, we found that *sog* is transcriptionally down regulated in the fatbody of tumour bearing animals (Figure 7 H-J). To determine if Sog downregulation contributes to cachexia progression, we overexpressed Sog in the fatbody, muscle or tumour of tumour-bearing animals. In all cases, tumour induced muscle detachment was significantly rescued (Figure 7 K-S). We also found that overexpression of Sog in the fatbody rescued Nidogen accumulation in the fatbody (Figure 7 T-V), indicating that the downregulation of *sog* drives cachexia by triggering fatbody ECM accumulation. This likely occurs via the activation of TGF-β signalling, which we have shown causes fatbody ECM accumulation (Figure 3, (Lodge et al., 2021)). Together, our data show that Sog is a critical regulator of cachexia.

## Discussion

It has been shown that cachexia is mediated by humoral factors secreted from the tumour, and the complete removal of cachexia-associated tumours can reverse cachexia (Ni et al., 2012). However, in late stage/ metastatic cancer patients, the removal of the tumour is often not an option. In this study, we demonstrate, using a *Drosophila* tumour model, that targeting signalling deregulations in the peripheral tissues can be a novel strategy to treat cachexia even when tumours are present, and the signalling status of peripheral tissues can serve as diagnostic biomarker for cachexia. We show that the insulin and TGF-β signalling pathways are deregulated in the peripheral tissues in tumour bearing animals, their disruption can recapitulate some aspects of tissue wasting, and their modulation can rescue tissue wasting in the presence of a tumour.

We have identified the fatbody as a central node in our cachexia model. In both fly and mouse cachexia models, the breakdown of fat precedes that of the muscles (Das et al., 2011; Lodge et al., 2021). Here, we find that tumour secreted proteins ImpL2 and Gbb converge in the fatbody to induce the activation of TGF-β signalling (Figure 7W). TGF-β activation then drives an aberrant accumulation of ECM at the inter-adipocyte junctions in the fatbody, which likely prevents ECM secretion and subsequent transport to the muscle. We found that the modulation of ECM secretion via SPARC or Rab10 can ameliorate the cachexia phenotype. It was previously reported that plasma membrane (PM) overgrowth induced by increased inflammation can cause pericellular collagen accumulation in the fatbody (Zang et al., 2015). In the cachectic fatbody, we have not observed PM thickening (data not shown), therefore, it is likely this occurs in cachexia via a different mechanism. Aberrant ECM accumulation in the adipose tissue has been observed in human cachexia patients, and this accumulation has been associated with elevated levels of TGF-β signalling (Alves et al., 2017). Furthermore, reduced ECM levels have been reported in the muscles in a rat cachexia model (Moraes et al., 2017). Thus, altered ECM localisation appears to be a conserved mechanism in cancer cachexia.

In this study, we show that insulin signalling (regulated by tumour derived ImpL2) directly affects muscle translation rates and atrophy. This, together with fatbody TGF-β activation/ ECM accumulation, appears to contribute toward the regulation of muscle integrity in the context of cachexia (Figure 7W). Additional tumour-secreted signals likely exist. Whether these signals also act through the fatbody (through regulation of ECM production or via alternative mechanisms), or act directly on the muscle or even via other target tissues, remains to be addressed. Studying these signals, their downstream signalling, as well as identifying additional target tissues where the signalling is relevant, remain important areas of future research.

The insulin signalling pathway plays many important roles in the regulation of growth and metabolism. TGF-β signalling is best known for roles in development, growth and differentiation. More recently, these pathways have been shown to intersect in metabolic contexts. In the fatbody, it has been shown under high fat diet induced obesity, that fatbody derived Gbb can cause insulin resistance via negative regulation of the insulin signalling pathway (Hong et al., 2016). Furthermore, in both *Drosophila* and *C. elegans,* it has been shown that BMP signalling can regulate Ilps to modulate lipid homeostasis (Ballard et al., 2010; Clark et al., 2021). In the developing muscle, it was recently shown that the Activin branch of the TGF-β signalling pathway positively regulates the insulin signalling pathway via the regulation of a structural protein, MHC (Kim and O’Connor, 2021). In contrast to these studies, we show that, in the fatbody, insulin signalling regulates TGF-β signalling by its modulation at the extracellular level via the secreted BMP inhibitor Sog. Interestingly, it has been shown in Zebrafish that IGF signalling can influence the expression of BMPs via Chordin (Sog homolog) during embryonic pattering, indicating that this relationship is conserved across distantly related species (Eivers et al., 2004). Although it was previously shown that the insulin signalling effector FOXO can directly bind to the promoter of Sog in adult flies (Birnbaum et al., 2019), this does not seem to be the case in the larval fatbody as FOXO overexpression did not significantly alter fatbody pMad levels (Figure 6). Instead, our data suggest that insulin signalling regulates *sog* expression via TOR and S6K. In support of this, TOR has been shown to influence other transcriptional regulators such as Raptor and Yorkie, thus it may regulate *sog* expression by modulating one of these transcriptional regulators (Parker and Struhl, 2015; Tiebe et al., 2015)

Sog has previously been shown to be important in embryonic patterning as well as at the developing cross-veins in the pupal wing. Our study also demonstrates that the inhibition of extracellular BMP ligands via Sog is a novel link between insulin and TGF-β signalling in the fatbody. We show that Sog is highly relevant in the context of cachexia, as it is found to be highly downregulated in the haemolymph of tumour-bearing animals. In addition, tissue specific overexpression of *sog* in the tumour, fatbody and muscle significantly rescued muscle integrity in the tumour bearing animals. Therefore, it appears that Sog levels in the tumour, as well as in peripheral tissues, play a role in mediating cachexia. As RNAseq data from the peripheral tissues of cachexia patients are not readily available through public data bases, we instead examined cBioPortal data which assesses gene expression in the tumour (9409 samples in 25 pan-cancer studies) (Cerami et al., 2012; Gao et al., 2013). We found that, in the tumour samples, there is a weak negative correlation between BMP7 expression (52% homology with gbb) and CHORDIN (CHRD, 40% homology with Sog) (Figure S7). This suggests that BMP and its regulators are expressed in many types of cancers, and that they may regulate each other’s expression. As such, CHRD may also be relevant for the progression of cachexia in humans, thus it would be interesting to explore whether circulating CHRD can serve as a potential biomarker for the diagnosis of cachexia. Together, our data suggest that TGF-β and insulin signalling crosstalk is relevant in the context of cancer cachexia. Our data also reveal potential therapeutic avenues to treat cachexia in the clinic when resection of the tumour is not an option in late stage/metastatic patients. These include targeting BMP antagonists such as Sog/CHRD, inhibition of TGF-β signalling, activation of insulin signalling, or enhancement of ECM secretion in the adipose tissue.

## Acknowledgements

We are grateful to Kieran Harvey, Helena Richardson, Gary Hime, Anne Holz, Travis Johnson, Christen Mirth and Ana Traven for generous sharing of fly stocks, yeast stock and antibodies. We would like to thank Bloomington Drosophila Stock Center, Vienna Drosophila Resource Center and Developmental Studies Hybridoma Bank for fly stocks and antibodies. We would like to also thank OZDros for *Drosophila* quarantine, Peter MacCallum Cancer Institute Microscopy Core for technical assistance. We are grateful to Kellie Veen and Khanh Phuong Nguyen for assistance with graphics production. LYC’s laboratory is supported by funding from the NHMRC Ideas Grant (APP2011289), ARC and the Peter MacCallum Cancer Foundation.

## Author contribution

D.B, S.G, C.D, E.C and L.Y.C conducted the experiments; B.J.S and B.M.S provided key reagents, D.B and L.Y.C wrote the paper.

---

**Figure S1:**
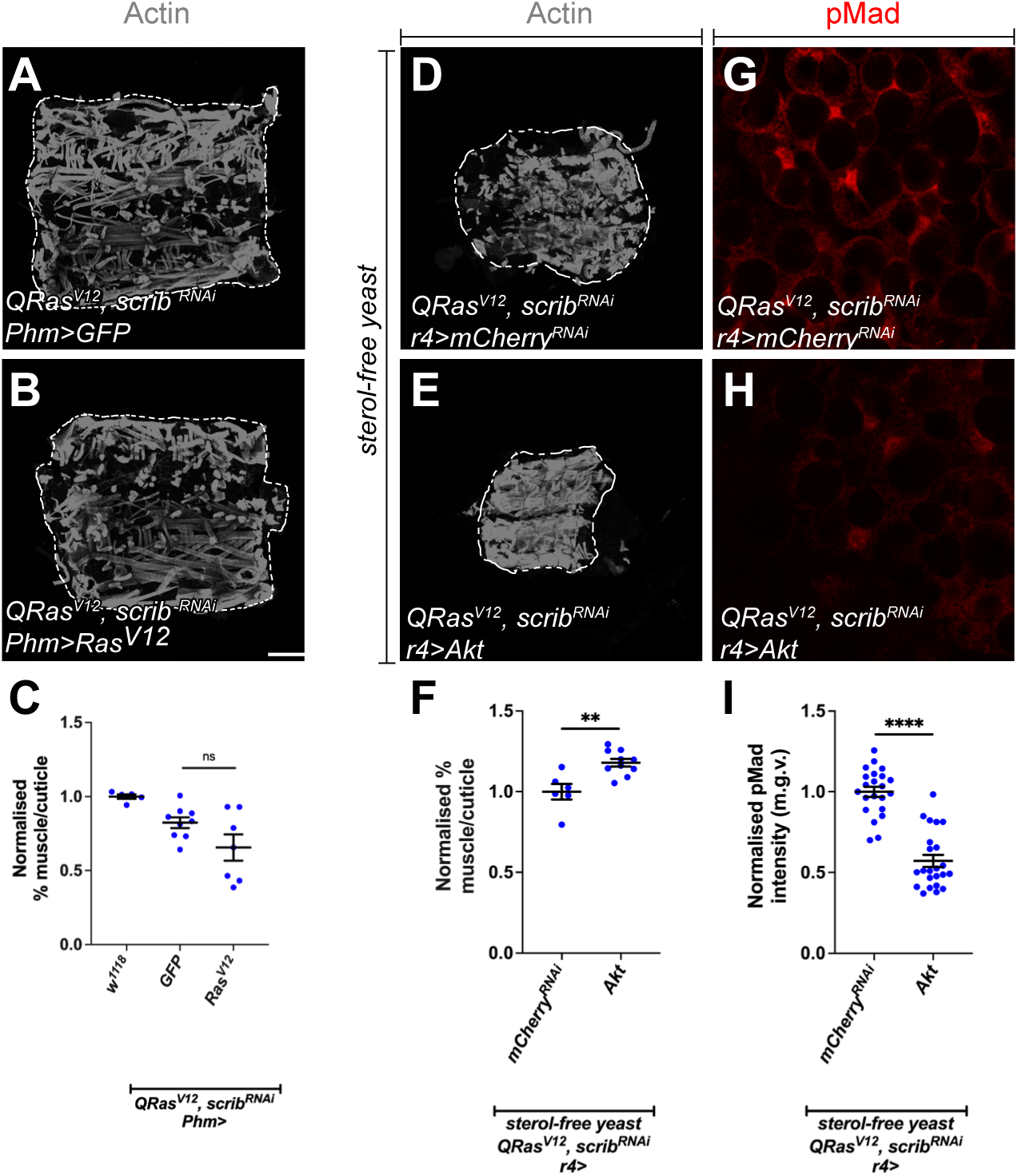
Systemic ecdysone levels does not affect muscle detachment. (A-B) Muscle fillets from animals where *GFP* or *Ras^V12^*was expressed in the prothoracic gland (*Phm-GAL4*) of *QRas^V12^scrib^RNAi^* tumour-bearing animals, exhibited similar amount of muscle detachment. (C) Quantification of muscle detachment in A-B. *w^1118^* (n=5), *GFP* (n=9), *Ras^v12^* (n=7). (D-E) Muscle fillets stained with phalloidin (Actin) from animals where *mCherry^RNAi^* or *UAS-Akt* was specifically expressed in the fatbody (*r4-GAL4*) of *QRas^V12^scrib^RNAi^* tumour-bearing animals fed on a sterol-free diet. (F) Quantification of muscle detachment in D-E. *mCherry^RNAi^* (n=6), *UAS-Akt* (n=10) (G-I) Fatbody from animals expressing *mCherry^RNAi^* or UAS-*Akt* under the control of *r4-GAL4* in *QRas^V12^scrib^RNAi^*tumour-bearing animals fed on a sterol-free diet stained with pMad. (J) Quantification of normalized fatbody pMad intensity in G-I*. mCherry ^RNAi^* (n=21), *UAS-Akt* (n=23). Scale bar is 500µm for muscle fillets and 50µm for fatbody staining.

**Figure S2:**
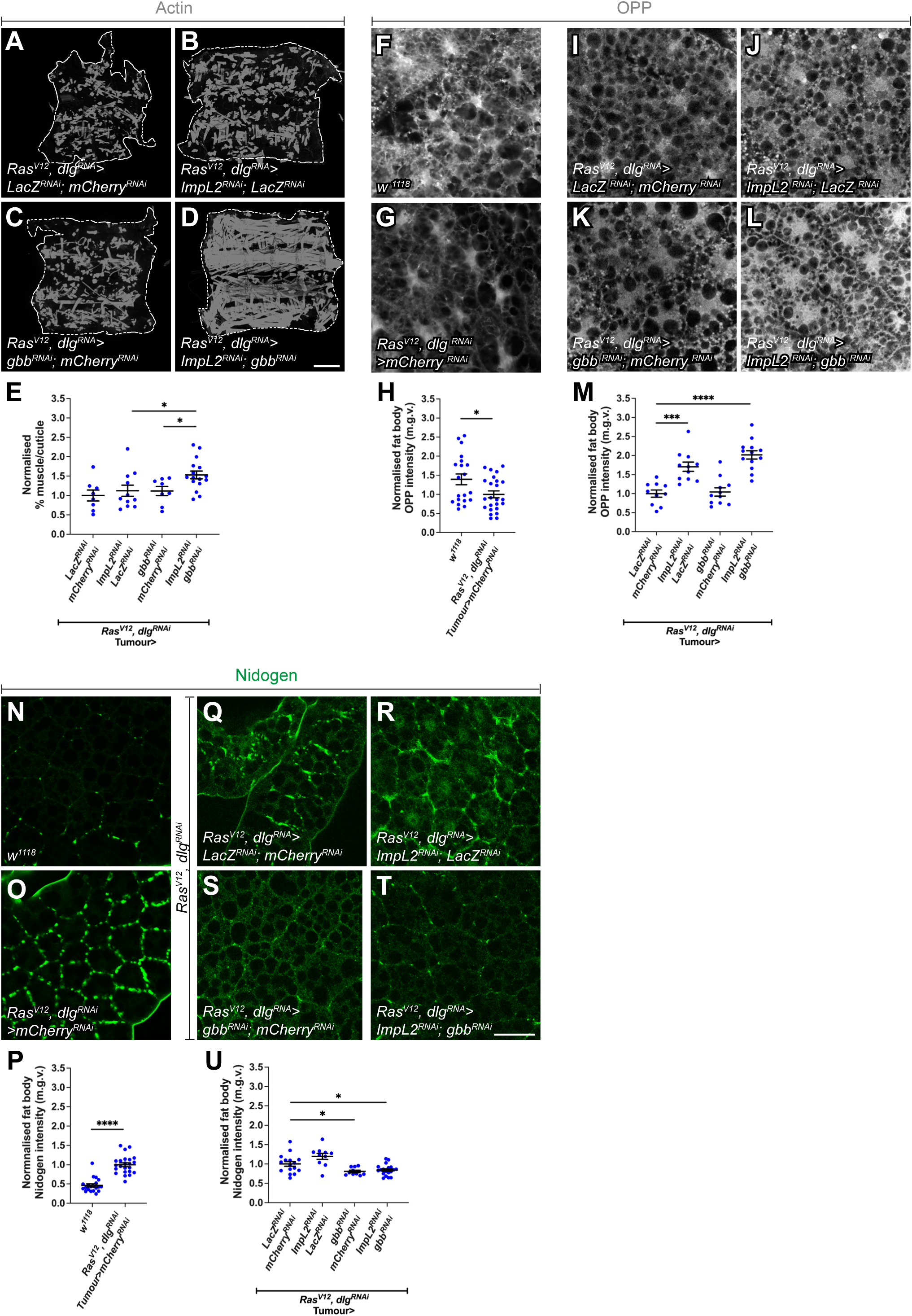
Knockdown of tumour derived Gbb and ImpL2 rescues muscle OPP and ECM protein Nidogen. (A) Tumour qPCRs showing mRNA expression levels of *gbb* in *Ras^V12^dlg^RNAi^* larvae expressing *mCherry^RNAi^* or *ImpL2^RNAi^*in the tumour (n=3). (B) Quantification of normalised muscle OPP intensity in *w^1118^* and *Ras^V12^dlg^RNAi^* tumour-bearing animals, where *mcherry^RNAi^* is specifically expressed in the tumour. *w^1118^* (n=15) and *Ras^V12^dlg^RNAi^*(n=15) (C-D) OPP staining detecting protein translation in the muscles of *Ras^V12^dlg^RNAi^* tumour-bearing animals that express *lacZ^RNAi^; mCherry^RNAi^* or *ImpL2^RNAi^; gbb^RNAi^*. (E) Quantification of OPP in C-D. *lacZ^RNAi^; mCherry^RNAi^* (n=8), *ImpL2^RNAi^; gbb^RNAi^* (n=10). (F) Quantification of normalised muscle Nidogen intensity in *w^1118^* and *Ras^V12^dlg^RNAi^* tumour-bearing animals, where *mcherry^RNAi^* is specifically expressed in the tumour. *w^1118^* (n=14) and *Ras^V12^dlg^RNAi^*(n=13) (G-H) Nidogen staining detecting ECM localisation in the muscles of *Ras^V12^dlg^RNAi^* tumour-bearing animals that express *lacZ^RNAi^; mCherry^RNAi^* or *ImpL2^RNAi^;gbb^RNAi^*. (I) Quantification of Nidogen in G-H. *lacZ^RNAi^; mCherry^RNAi^* (n=7), *ImpL2^RNAi^; gbb^RNAi^* (n=15). Scale bar is 50µm.

**Figure S3:**
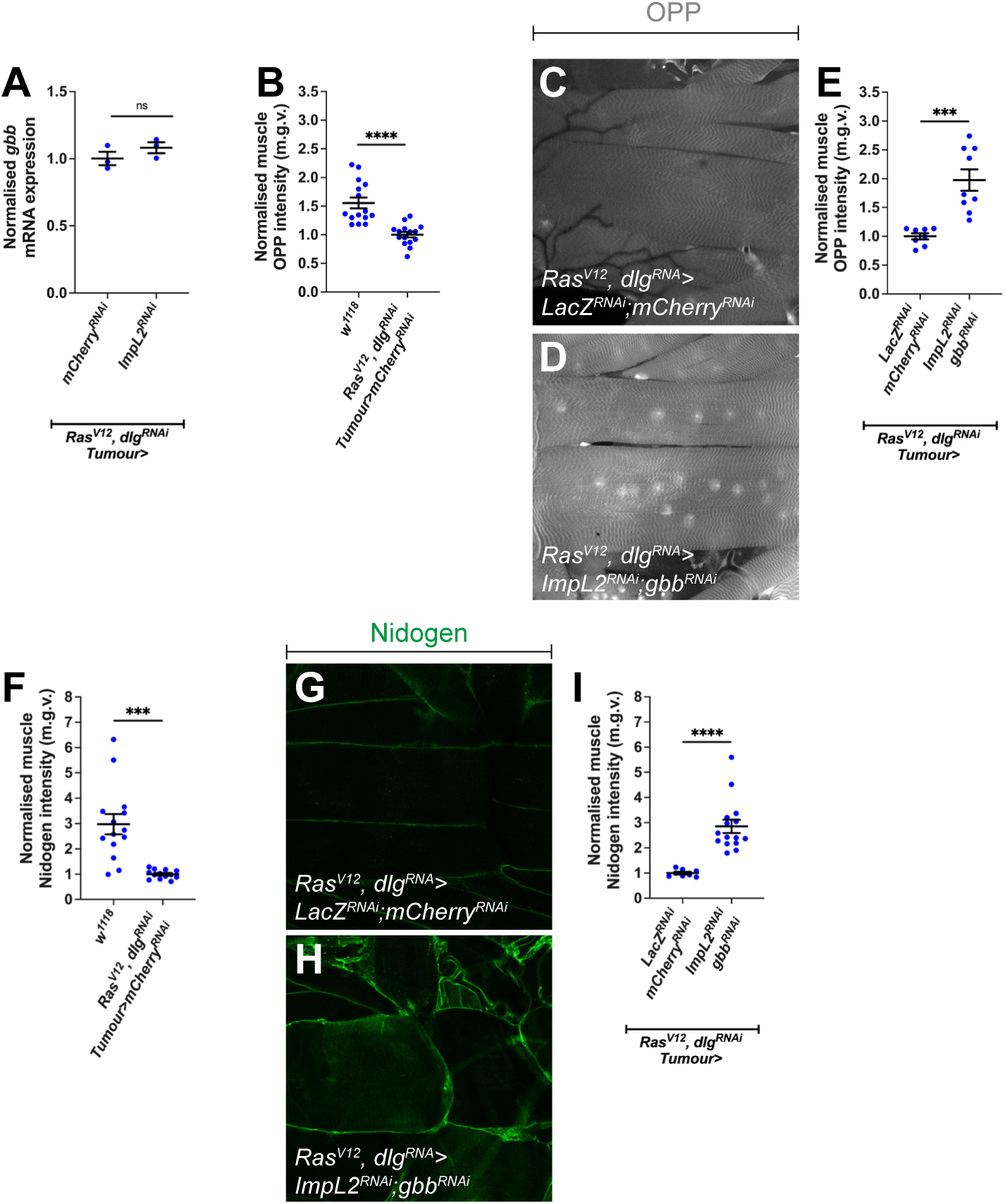
Tumour secreted Gbb and ImpL2 rescue muscle integrity additively. (A-D) Muscle fillets of *Ras^V12^dlg^RNAi^* tumour-bearing animals that express in the tumour *lacZ^RNAi^; mCherry^RNAi^* or *ImpL2^RNAi^; lacZ^RNAi^*, or *gbb^RNAi^; mCherry^RNAi^* or *ImpL2^RNAi^; gbb^RNAi^*. (E) Quantification of muscle detachment in A-D. *lacZ^RNAi^; mCherry ^RNAi^* (n=8), *ImpL2^RNAi^; lacZ^RNAi^*(n=11), *gbb^RNAi^; mCherry^RNAi^* (n=8), *ImpL2^RNAi^; gbb^RNAi^* (n=16). (F-G) OPP staining detecting protein translation in the fatbody of *w^1118^* and *Ras^V12^dlg^RNAi^* tumour-bearing animals. (H) Quantification of OPP in F-G. *w^1118^* (n=19) *Ras^V12^dlg^RNAi^*(n=26) (I-L) OPP staining detecting protein translation in the fatbody of *Ras^V12^dlg^RNAi^* tumour-bearing animals, that express in the tumour *lacZ^RNAi^; mCherry^RNAi^* or *ImpL2^RNAi^; lacZ^RNAi^*, or *gbb^RNAi^; mCherry^RNAi^*or *ImpL2^RNAi^; gbb^RNAi^*. (M) Quantification of OPP in I-L. *lacZ^RNAi^; mCherry^RNAi^* (n=10), *ImpL2^RNAi^; lacZ^RNAi^* (n=11), *gbb^RNAi^; mCherry^RNAi^* (n=11), *ImpL2^RNAi^; gbb^RNAi^* (n=13). (N-O) Nidogen staining detecting ECM localisation in the fatbody of *w^1118^* and *Ras^V12^dlg^RNAi^* animals. (P) Quantification of nidogen staining in N-O. *w^1118^* (n=23) *Ras^V12^dlg^RNAi^*(n=20) (R-U) *Ras^V12^dlg^RNAi^* tumour-bearing animals that express *lacZ^RNAi^; mCherry^RNAi^* or *ImpL2^RNAi^; lacZ^RNAi^*, or *gbb^RNAi^; mCherry^RNAi^* or *ImpL2^RNAi^; gbb^RNAi^*. (V) Quantification of Nidogen in *lacZ^RNAi^; mCherry^RNAi^* (n=16), *ImpL2^RNAi^; lacZ^RNAi^* (n=10), *gbb^RNAi^*

**Figure S4:**
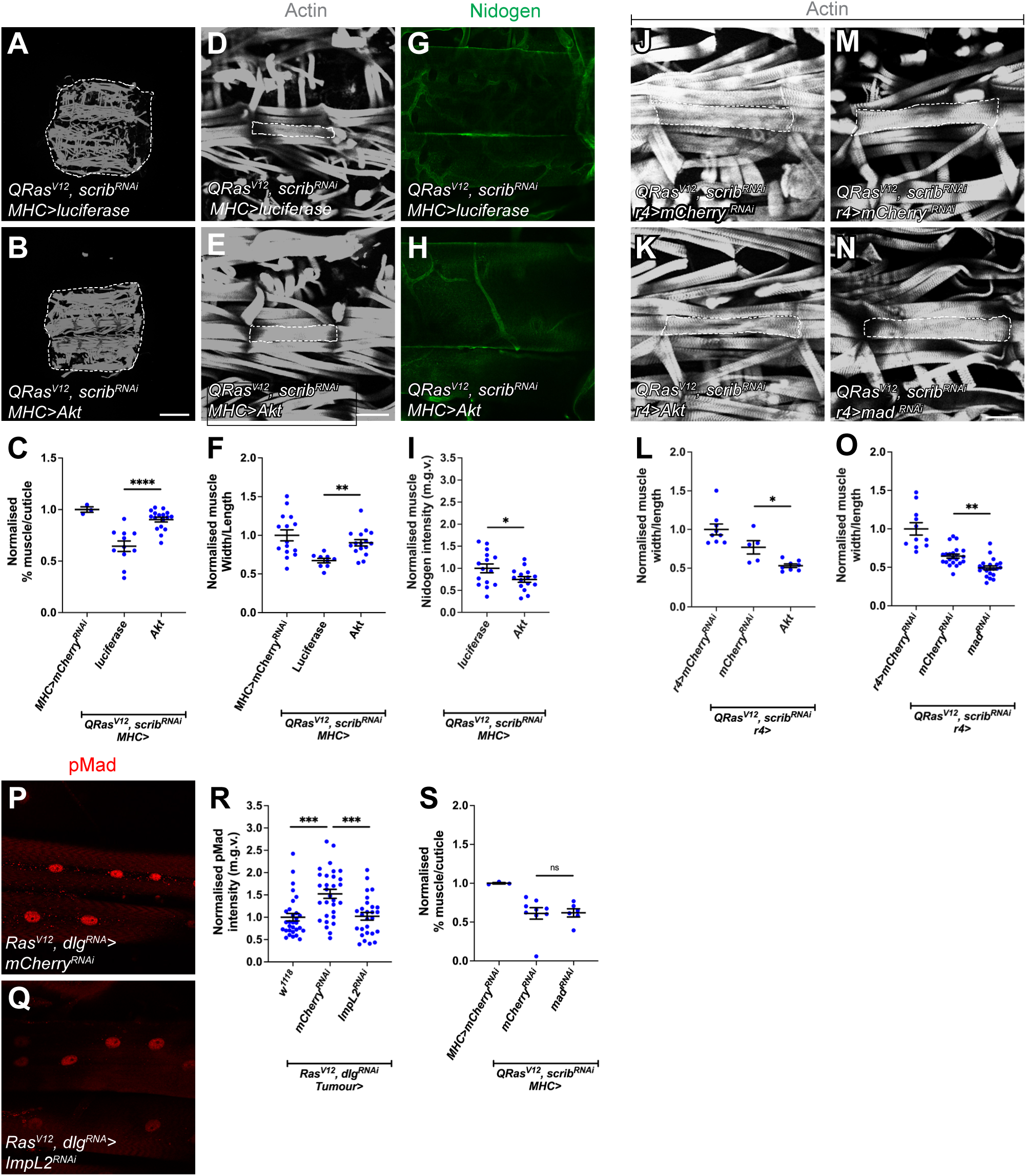
Muscle overexpression of Akt (but not fatbody Akt or mad RNAi overexpression) rescues atrophy in cachectic animals, and TGF-ß signalling in the muscle is responsive to modulation in tumour insulin signalling but is not required for muscle integrity in cachexia. (A-B) Muscle fillets stained with phalloidin (Actin) where *luciferase* or *Akt* was expressed under the control of *MHC-GAL4* in *QRas^V12^scrib^RNAi^* tumour-bearing animals. (C) Quantification of normalized muscle detachment in A-B. *w^1118^* (n=3), *UAS-luciferase* (n=10), *Akt* (n=17). (D-E) Muscle segment (outlined) where *luciferase* or *Akt* was expressed under the control of *MHC-GAL4* in *QRas^V12^scrib^RNAi^* tumour-bearing animals. (F) Quantification of normalized muscle width/length in D-E. *w^1118^* (n=15), *luciferase* (n=9), *Akt* (n=16). (G-H) Muscle nidogen staining where *luciferase* or *Akt* was expressed under the control of *MHC-GAL4* in *QRas^V12^scrib^RNAi^* tumour-bearing animals. (I) Quantification of normalized muscle nidogen staining in G-H. *luciferase* (n=16), *Akt* (n=15). (J-K) Muscle segment from *QRas^V12^scrib^RNAi^* tumour-bearing animals, where *mCherry^RNAi^* or *Akt* was expressed in the fatbody (*r4-GAL4*). (L) Quantification of normalized muscle width/length in D-E. *r4>mCherry^RNAi^*(n=9), *QRas^V12^scrib^RNAi^;r4> mCherry^RNAi^* (n=6), *QRas^V12^scrib^RNAi^;r4>Akt* (n=21). (M-N) Muscle segment from *QRas^V12^scrib^RNAi^* tumour-bearing animals, where *mCherry^RNAi^* or *mad^RNAi^* was expressed in the fatbody (*r4-GAL4*). (O) Quantification of normalized muscle width/length in D-E. *r4>mCherry^RNAi^*(n=11), *QRas^V12^scrib^RNAi^; r4> mCherry^RNAi^* (n=18), *QRas^V12^scrib^RNAi^; r4> mad^RNAi^* (n=15). (P-Q) Muscle from *Ras^V12^Dlg^RNAi^* tumour-bearing animals expressing either *mCherry ^RNAi^* or *ImpL2^RNAi^* in the tumour, where TGF-ß signaling activation is indicated by pMad staining. (R) Quantification of normalized muscle pMad intensity in (P-Q)*. w^1118^* (n=30)*, mCherry^RNAi^* (n=30), *ImpL2^RNAi^* (n=26). (S) Quantification of normalized muscle detachment where *MHC>mCherry^RNAi^* (n=9) or *mCherry^RNAi^* (n=9) or *lacZ^RNAi^; mad^RNAi^*(n=6) was expressed under the control of *MHC-GAL4* in *QRas^V12^scrib^RNAi^*tumour-bearing animals. Scale bar is 500µm for muscle fillets, 100µm for muscle segment atrophy measurements and 50µm for muscle Nidogen and pMad staining.

**Figure S5:**
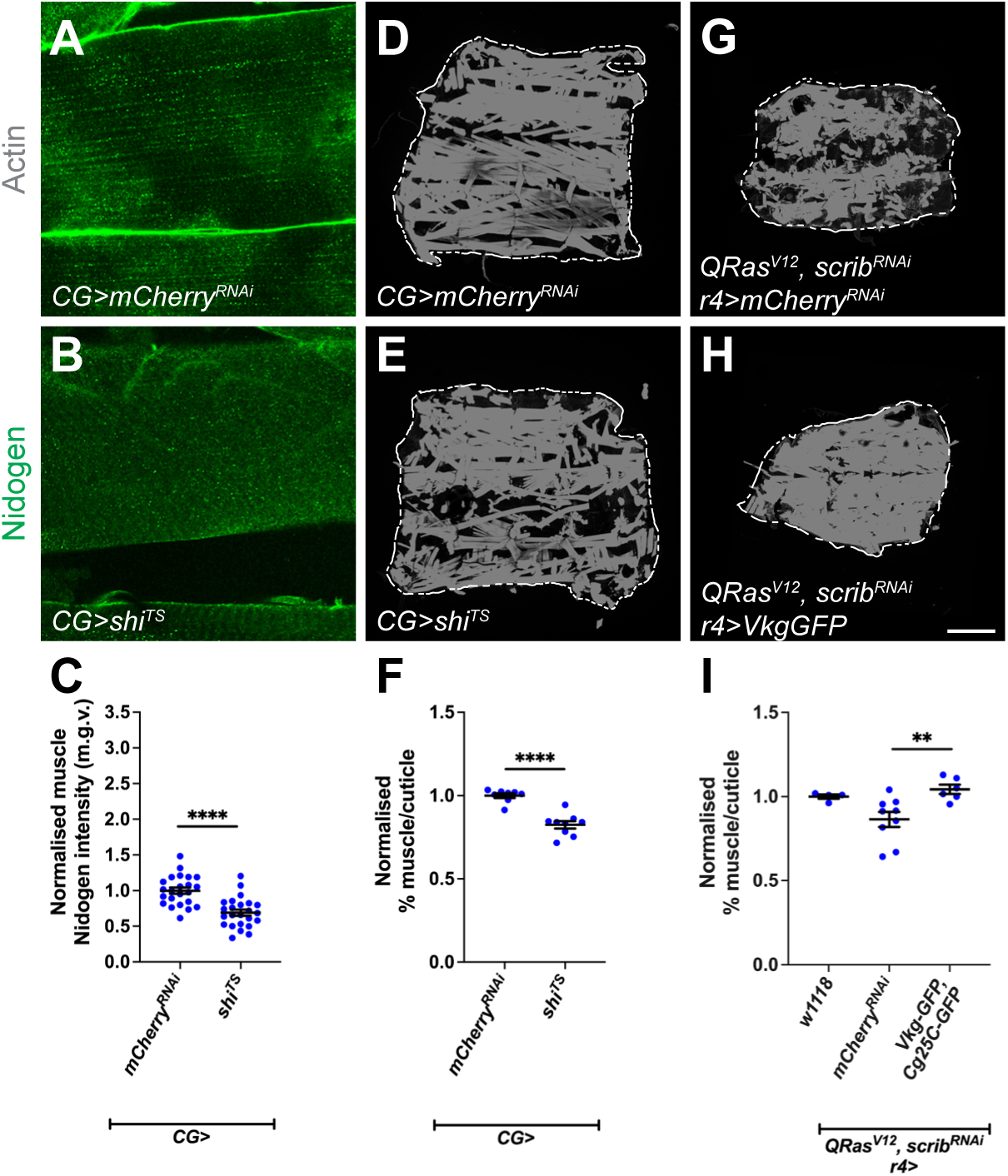
Inhibition of fatbody endocytosis causes muscle detachment, increasing Vkg expression in the fatbody improves muscle integrity in tumour bearing animals. (A-B) Nidogen staining in muscles of animals where *mCherry^RNAi^* or *shi^TS^* was expressed in the fatbody via *CG-GAL4*. (C) Quantification of muscle nidogen staining in A-B. *mCherry^RNAi^* (n=23), *shi^TS^* (n=24). (D-E) Muscle fillets stained with phalloidin (Actin) from animals where *mCherry^RNAi^* or *shi^TS^* was expressed in the fatbody via *CG-GAL4*. (F) Quantification of normalized muscle detachment in D-E. *mCherry^RNAi^* (n=8), *shi^TS^* (n=9). (G-H) Muscle fillets stained with phalloidin (Actin) from *QRas^V12^scrib^RNAi^*tumour-bearing animals, where *mCherry^RNAi^* or *UAS-VkgGFP;UAS-Cg25C-GFP* was expressed in the fatbody (*r4-GAL4*). Day 6 muscles used here. (I) Quantification of normalized muscle detachment in G-H. *w^1118^* (n=4), *QRas^V12^scrib^RNAi^;r4> mCherry^RNAi^* (n=9), *QRas^V12^scrib^RNAi^;r4>UAS-VkgGFP;UAS-Cg25C-GFP* (n=6). Scale bar is 500µm for muscle fillet, 100µm for muscle segment atrophy measurements, 50µm for muscle nidogen staining and 50µm for fatbody staining. Graphs are represented as Mean ± SEM, n= the number of samples. *(*) p<0.05 (**) p< 0.01, (***) p< 0.001, (****) p < 0.0001, (ns) p>0.05*

**Figure S6:**
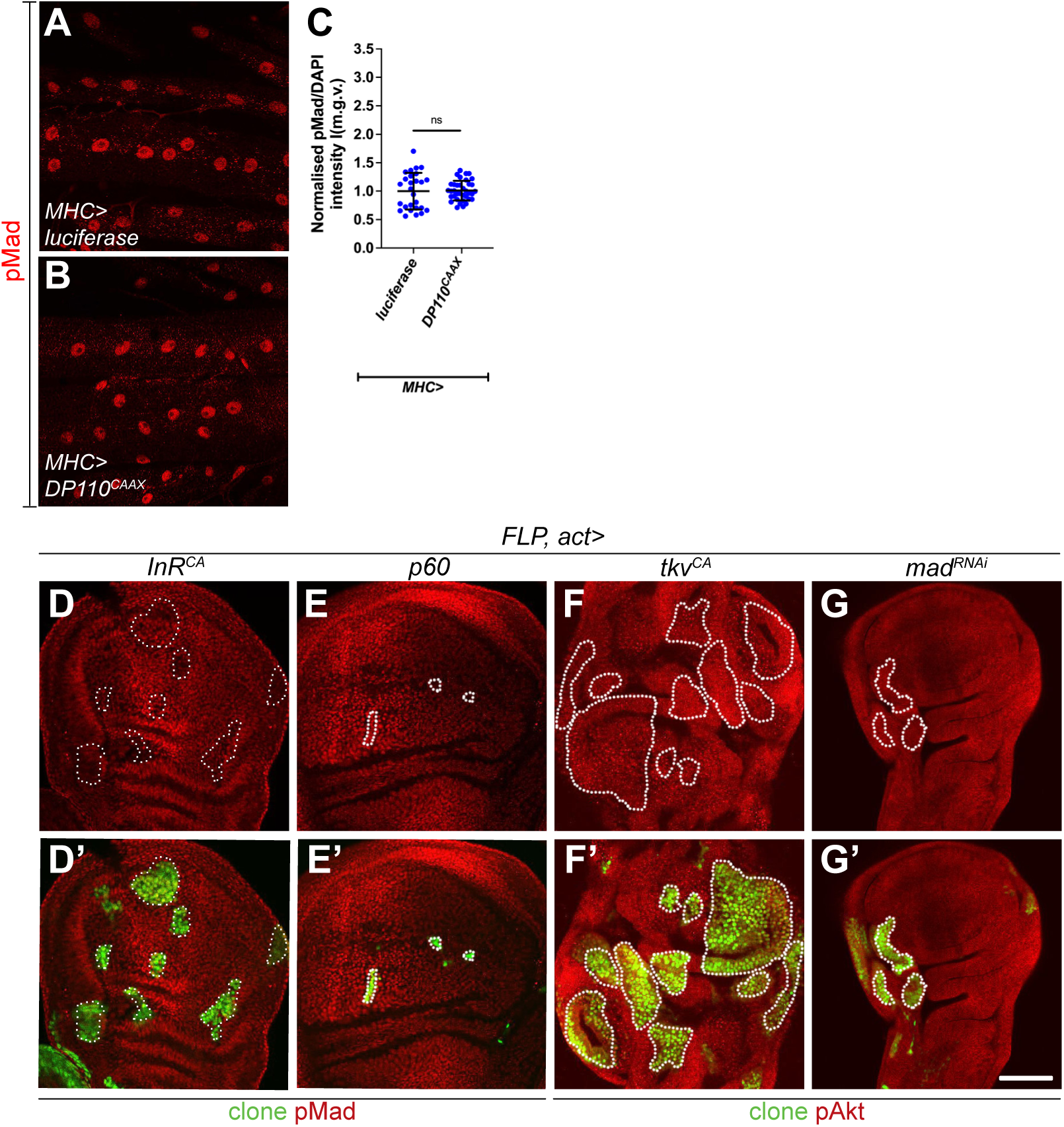
Modulation of TGF-ß signalling does not affect insulin signalling in the muscle or wing imaginal discs. (A-B) pMad staining in muscle fillets from larvae expressing *luciferase* or *DP110^CAAX^* with *MHC-GAL4*. (C) Quantification of pMad staining in A-B. *luciferase* (n=27), *DP110^CAAX^* (n=36). (D-D’) Heatshock induced flip-out clones (marked with GFP, circled with dotted lines) of *InR^CA^* in the wing imaginal discs, readout of TGF-ß signalling levels indicated by pMad is not altered in the clones vs. surrounding tissues. (E-E’) Heatshock induced flip-out clones (marked with GFP, circled with dotted lines) of *p60* in the wing imaginal discs, readout of TGF-ß signalling levels indicated by pMad is not altered in the clones vs. surrounding tissues. (F-F’) Heatshock induced flip-out clones (marked with GFP, circled with dotted lines) of *tkv^CA^* in the wing imaginal discs, readout of insulin signalling levels indicated by pAkt is not altered in the clones vs. surrounding tissues. (G-G’) Heatshock induced flip out clones (marked with GFP, circled with dotted lines) of *mad^RNAi^* in the wing imaginal discs, readout of insulin signalling levels indicated by pAkt is not altered in the clones vs. surrounding tissues. Scale bar is 50 µm.

**Figure S7.**
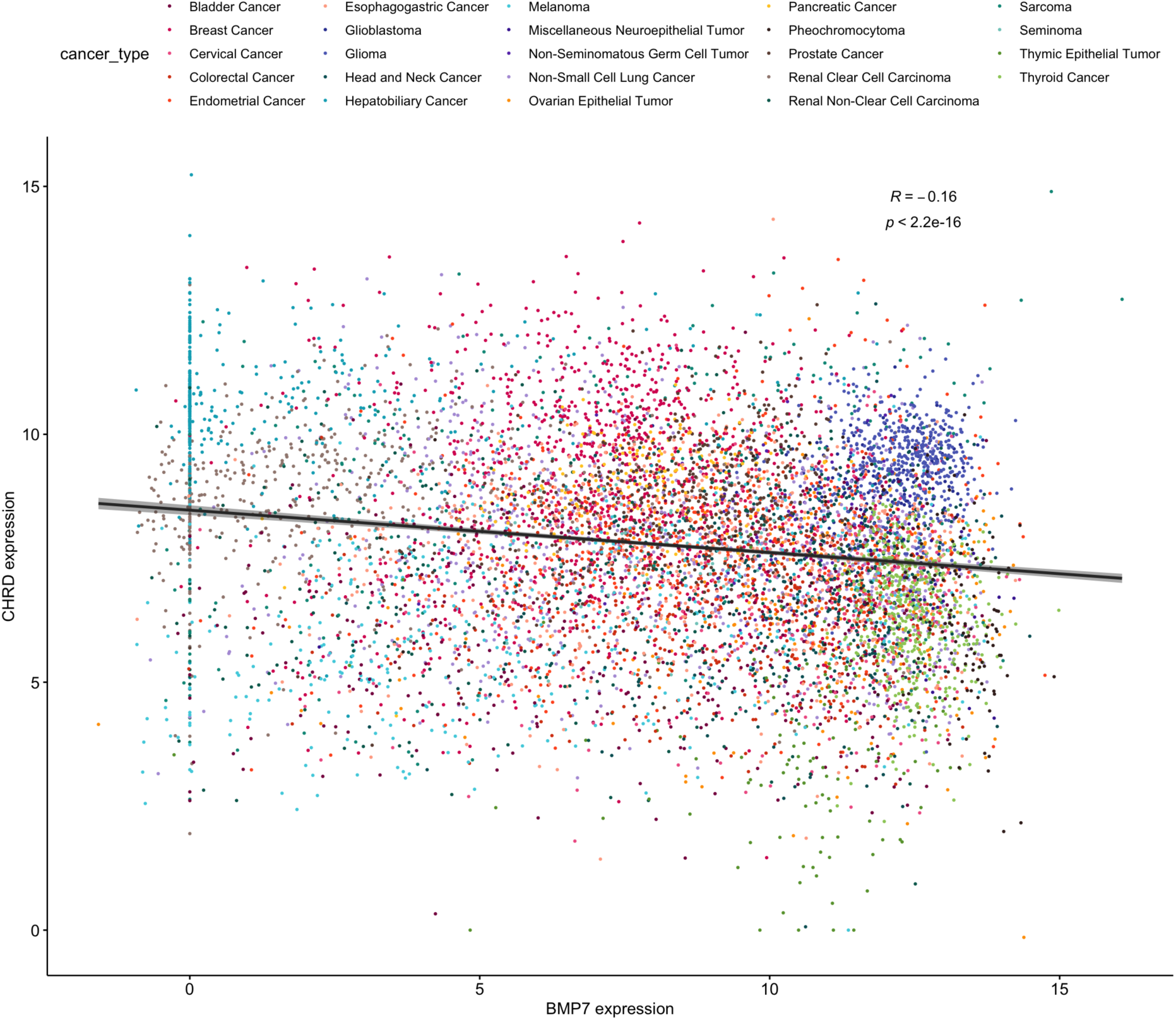
Pan-cancer expression of CHRD compared to BMP7. Data is RSEM batch normalized log2(value + 1) as per cBioPortal, points coloured by cancer type. Spearman correlation R and p-values shown. Regression line is shown in black.

## Methods

### *Drosophila* stocks and husbandary

The following stocks were used from the Bloomington *Drosophila* stock centre: *CG-GAL4* (BL7011), *Mef2-GAL4* (BL27391), *MHC-GAL4* (BL55133), *r4-GAL4* (BL33832), *UAS-4E-BP^CA^* (BL24854), *UAS-Akt* (BL8191), *UAS-Dp110^CAAX^* (BL25908), *UAS-Gbb^RNAi#2^* (BL34898), *UAS-InR^CA^* (BL8263), *UAS-luciferase* (BL64774), *UAS-mad^RNAi^* (BL31316), UAS-luciferase (BL64774), *UAS-mCherry^RNAi^* (BL35785), *UAS-S6K^CA^* (BL6914), *UAS-Shi^TS^* (BL44222), *UAS-tkv^CA^* (BL36536), *UAS-Sparc3* (Helena Richardson), ey-FLP1;act>CD2>GAL4,UAS-GFP (Lodge et al., 2021), sog-lacZ (BL10132). The following stocks were obtained from the Vienna Drosophila Resource Centre: *UAS-Gbb^RNAi^* (v330684), *UAS-Impl2^RNAi^* (v30931), *UAS-Sparc^RNAi^* (v16677). The following stock was obtained from the Kyoto stock centre: *Vkg-GFP* (110692). The following stocks were also used: *Ey-FLP1; QUAS-Ras^V12^, QUAS-scrib^RNAi^/ CyOQS; act>CD2>QF, UAS-RFP/TMBQS* (Lodge et al., 2021) *Ey-FLP1; UAS-Ras^V12^, UAS-dlg^RNAi^/ CyO, GAL80; act>CD2>GAL4, UAS-GFP* (Manent et al., 2017), *Phm-GAL4* (Christin Mirth), *UAS-FOXO* (Kieran Harvey), *UAS-GFP* (Kieran Harvey), *UAS-lacZ^RNAi^* (Kieran Harvey), *UAS-p60* (Cheng et al., 2011), *UAS-Ras^V12^* (Helena Richardson), *UAS-Sog-HA* (called Sog here, from Gary Hime), *UAS-TOR^DN^*(Zhang et al., 2006), *UAS-cg25C;UAS-Vkg* (Brain Stramer).

For the *r4>InR^CA^*; *mad^RNAi^* interaction experiment and analysis of atrophy in *QRas^V12^scrib^RNAi^*larvae carrying *r4>mad^RNAi^* larvae were reared at 25°C. For experiments with *CG*>*InR^CA^* larvae were reared at 18°C. For experiments with *CG>shi^TS^*, adults were allowed to lay for 24h at 25°C and progeny transferred to 31°C after an additional 48h. For all other experiments, adults were allowed to lay for 24h at 25°C and the progeny then moved to 29°C. Animals were dissected at wandering stage in non-tumour bearing animals(with the exception of the *CG>shi^TS^* experiments, conducted on day 7 animals), and for tumour-bearing animals, they were dissected on 7 days after egg lay or as indicated throughout.

### Immunostaining

For muscle staining, larvae were filleted as previously described (Dark et al., 2022; Lodge et al., 2021), fixed for 30min in PBS containing 4% formaldehyde and washed with PBS containing 0.3% Triton-X (PBST-0.3). Fatbody was fixed for 45min and washed with PBS containing 0.2% Triton-X (PBST-0.2). Wing discs were fixed for 20min and washed with PBS containing 0.2% Triton-X (PBST-0.2). Tissues were then stained as per the manufacturer’s specifications. Samples were mounted in glycerol and imaged on an Olympus FV3000 confocal microscope. Within a given experiment, all images were acquired using identical settings. Fat body samples were imaged the same day as mounting. To assay for translation in the muscle, we used Click-iT™ Plus OPP Alexa Fluor™ 488 Protein Synthesis Assay Kit (Thermo Fisher, #C10456). Primary antibodies used: ßgal (1:100, Promega), pMad (1:800, Cell Signalling, #9516, used for fatbody staining), pMad (1:100, abcam, ab52903, used for muscle staining) pAkt (Cell signalling, 1:100, #4060), Nidogen (gift of Anne Holtz, 1:500). Secondary donkey antibodies conjugated to Alexa 555 and Alexa 647 (Molecular Probes) were used at 1:200. DAPI (Molecular Probes) was used at 1:10000, Phalloidin (Molecular Probes) was used at 1:10000. Actin muscle stains were conducted on day 7 animals when dissected from *QRas^V12^scrib^RNAi^*animals except when raised at 25°C and on day 8 animals when dissected from *Ras^V12^dlg^RNAi^*. All other muscle and fatbody staining were conducted at day 6 when dissected from *QRas^V12^scrib^RNAi^* animals and at day 7 when dissected from *Ras^V12^dlg^RNAi^* animals (except when specified in the figure legend).

### Image analysis

All images were quantified using ImageJ. In wildtype fatbody and muscle samples pMad intensity was normalised to DAPI, pMad and DAPI levels were quantified by drawing a circle around the nucleus in the DAPI channel, and the mean grey value (m.g.v.) determined for pMad and DAPI channels. In tumour bearing fatbody, pMad, pAkt and OPP were not normalised to DAPI levels, as fatbody tissue penetrance is altered in the presence of tumour. Here, the ROI circle was drawn around the nucleus in the pMad, pAkt or OPP channel. To measure fluorescence intensity of Vkg-GFP and Ndg, a line was drawn around membrane of a single cell in the fatbody or along the edge of a single muscle segment in fillets on the z-plane where fluorescence was most intense, the line made to be five pixels wide, and m.g.v. determined along the line. %muscle/ cuticle was determined using ImageJ as previously described (Dark et al., 2022; Lodge et al., 2021). For muscle atrophy measurements, muscle length and width measurements were done on the VL3 muscle on tiled 10X images of individual fillets using FIJI software with the line selection tool.

### qPCR

For each biological replicate, three to five larvae were selected at day 6 (for fatbody samples) or day 7 (for tumour samples). Samples were dissected in cold PBS and snap frozen in liquid nitrogen. Frozen tissue was then lysed in 300μL of TRI Reagent (Invitrogen, #10296010) Total RNA was extracted using a Direct-zol RNA Microprep Kit (Zymo Research, #R2061). cDNA was obtained by reverse transcription of 0.5 μg of total RNA using ProtoScript II First Strand cDNA Synthesis Kit (NEB, #E6560S). Three independent biological replicates were prepared for each genotype. qPCR was performed with the Fast SYBR™ Green PCR Master Mix (Applied Biosystems, #4385612) on a LightCycler 480 (Roche). Gene expression was normalized to the geometric mean of the reference gene *rpl32*.

### TCGA tumour data analyses

mRNA expression data for CHRD and BMP7 was downloaded for 24 TCGA studies from cBioPortal on 3rd August 2022. A total of 9409 samples had RSEM data that had been batch normalised for both genes, and Spearman correlation was performed.

### Statistical analysis

Hemolymph analysis was done using Limma. All other statistical analyses were conducted using GraphPad Prism. At least three animals per genotype were used for all muscle and fatbody experiments. For staining intensity quantifications, individual data points represent fluorescence intensity of a single cell. For muscle integrity quantifications, individual data points represent a single larva. Data in Figure 4H was confirmed to be normally distributed using the Shapiro-Wilk test. For experiments with two genotypes or treatments, two-tailed unpaired student’s t-tests were used to test for significant differences. The Welch’s correction was applied in cases of unequal variances. For experiments with more than two genotypes, significant differences between specific genotypes were tested using a one-way ANOVA and a subsequent Šidák post-hoc test. The results for all post-hoc tests conducted in each analysis are shown in graphs. For all graphs, error bars represent SEM. p and adjusted-p values are reported as follows: p>0.05, ns (not significant); p<0.05, *; p<0.01, **; p<0.001, ***; p<0.0001, ****.

## References

Alves, M. J., Figuerêdo, R. G., Azevedo, F. F., Cavallaro, D. A., Neto, N. I. P., Lima, J. D. C., Matos-Neto, E., Radloff, K., Riccardi, D. M., Camargo, R. G., et al. (2017). Adipose tissue fibrosis in human cancer cachexia: the role of TGFβ pathway. BMC Cancer 17,.

Ballard, S. L., Jarolimova, J. and Wharton, K. A. (2010). Gbb/BMP signaling is required to maintain energy homeostasis in Drosophila. Dev Biol 337, 375–385.

Baracos, V. E., Martin, L., Korc, M., Guttridge, D. C. and Fearon, K. C. H. (2018). Cancer-associated cachexia. Nature Publishing Group 4, 1–18.

Biehs, B., François, V. and Bier, E. (1996). The Drosophila short gastrulation gene prevents Dpp from autoactivating and suppressing neurogenesis in the neuroectoderm. Genes Dev 10, 2922–2934.

Birnbaum, A., Wu, X., Tatar, M., Liu, N. and Bai, H. (2019). Age-Dependent Changes in Transcription Factor FOXO Targeting in Female Drosophila. Frontiers in Genetics 10,.

Brogiolo, W., Stocker, H., Ikeya, T., Rintelen, F., Fernandez, R. and Hafen, E. (2001). An evolutionarily conserved function of the Drosophila insulin receptor and insulin-like peptides in growth control. 11, 213–221.

Caldwell, P. E., Walkiewicz, M. and Stern, M. (2005). Ras Activity in the Drosophila Prothoracic Gland Regulates Body Size and Developmental Rate via Ecdysone Release. Current Biology 15, 1785–1795.

Cerami, E., Gao, J., Dogrusoz, U., Gross, B. E., Sumer, S. O., Aksoy, B. A., Jacobsen, A., Byrne, C. J., Heuer, M. L., Larsson, E., et al. (2012). The cBio cancer genomics portal: an open platform for exploring multidimensional cancer genomics data. Cancer Discov 2, 401–404.

Cheng, L. Y., Bailey, A. P., Leevers, S. J., Ragan, T. J., Driscoll, P. C. and Gould, A. P. (2011). Anaplastic Lymphoma Kinase Spares Organ Growth duringNutrient Restriction in Drosophila. Cell 146, 435–447.

Clark, J. F., Ciccarelli, E. J., Kayastha, P., Ranepura, G., Yamamoto, K. K., Hasan, M. S., Madaan, U., Meléndez, A. and Savage-Dunn, C. (2021). BMP pathway regulation of insulin signaling components promotes lipid storage in Caenorhabditis elegans. PLOS Genetics 17, e1009836.

Colombani, J., Bianchini, L., Layalle, S., Pondeville, E., Dauphin-Villemant, C., Antoniewski, C., Carré, C., Noselli, S. and Léopold, P. (2005). Antagonistic actions of ecdysone and insulins determine final size in Drosophila. Science 310, 667–670.

Dai, J., Ma, M., Feng, Z. and Pastor-Pareja, J. C. (2017). Inter-adipocyte Adhesion and Signaling by Collagen IV Intercellular Concentrations in Drosophila. CURBIO 27, 2729–2740.e4.

Dai, J., Estrada, B., Jacobs, S., Sánchez-Sánchez, B. J., Tang, J., Ma, M., Magadán-Corpas, P., Pastor-Pareja, J. C. and Martín-Bermudo, M. D. (2018). Dissection of Nidogen function in Drosophila reveals tissue-specific mechanisms of basement membrane assembly. PLoS Genet 14, e1007483.

Dark, C., Cheung, S. and Cheng, L. Y. (2022). Analyzing cachectic phenotypes in the muscle and fat body of Drosophila larvae. STAR Protoc 3, 101230.

Das, S. K., Eder, S., Schauer, S., Diwoky, C., Temmel, H., Guertl, B., Gorkiewicz, G., Tamilarasan, K. P., Kumari, P., Trauner, M., et al. (2011). Adipose triglyceride lipase contributes to cancer-associated cachexia. Science 333, 233–238.

Ding, G., Xiang, X., Hu, Y., Xiao, G., Chen, Y., Binari, R., Comjean, A., Li, J., Rushworth, E., Fu, Z., et al. (2021). Coordination of tumor growth and host wasting by tumor-derived Upd3. Cell Rep 36, 109553.

Dong, J., Yu, J., Li, Z., Gao, S., Wang, H., Yang, S., Wu, L., Lan, C., Zhao, T., Gao, C., et al. (2021). Serum insulin-like growth factor binding protein 2 levels as biomarker for pancreatic ductal adenocarcinoma-associated malnutrition and muscle wasting. Journal of Cachexia, Sarcopenia and Muscle 12, 704–716.

Eivers, E., McCarthy, K., Glynn, C., Nolan, C. M. and Byrnes, L. (2004). Insulin-like growth factor (IGF) signalling is required for early dorso-anterior development of the zebrafish embryo. Int J Dev Biol 48, 1131–1140.

Fearon, K., Arends, J. and Baracos, V. (2013). Understanding the mechanisms and treatment options in cancer cachexia. Nature reviews. Clinical oncology 10, 90–99.

Figueroa-Clarevega, A. and Bilder, D. (2015). Malignant Drosophila Tumors Interrupt Insulin Signaling to Induce Cachexia-like Wasting. Developmental Cell 33, 47–55.

Gao, J., Aksoy, B. A., Dogrusoz, U., Dresdner, G., Gross, B., Sumer, S. O., Sun, Y., Jacobsen, A., Sinha, R., Larsson, E., et al. (2013). Integrative analysis of complex cancer genomics and clinical profiles using the cBioPortal. Sci Signal 6, pl1.

Gesualdi, S. C. and Haerry, T. E. (2007). Distinct signaling of Drosophila Activin/TGF-beta family members. Fly (Austin) 1, 212–221.

Giannakou, M. E. and Partridge, L. (2007). Role of insulin-like signalling in Drosophila lifespan. Trends in Biochemical Sciences 32, 180–188.

Graca, F. A., Rai, M., Hunt, L. C., Stephan, A., Wang, Y.-D., Gordon, B., Wang, R., Quarato, G., Xu, B., Fan, Y., et al. (2022). The myokine Fibcd1 is an endogenous determinant of myofiber size and mitigates cancer-induced myofiber atrophy. Nat Commun 13, 2370.

Hodgson, J. A., Parvy, J.-P., Yu, Y., Vidal, M. and Cordero, J. B. (2021). Drosophila Larval Models of Invasive Tumorigenesis for In Vivo Studies on Tumour/Peripheral Host Tissue Interactions during Cancer Cachexia. Int J Mol Sci 22, 8317.

Honegger, B., Galic, M., Köhler, K., Wittwer, F., Brogiolo, W., Hafen, E. and Stocker, H. (2008). Imp-L2, a putative homolog of vertebrate IGF-binding protein 7, counteracts insulin signaling in Drosophila and is essential for starvation resistance. J Biol 7, 10.

Hong, S.-H., Kang, M., Lee, K.-S. and Yu, K. (2016). High fat diet-induced TGF-β/Gbb signaling provokes insulin resistance through the tribbles expression. Sci Rep 6, 30265.

Isabella, A. J. and Horne-Badovinac, S. (2016). Rab10-Mediated Secretion Synergizes with Tissue Movement to Build a Polarized Basement Membrane Architecture for Organ Morphogenesis. Developmental Cell 38, 47–60.

Khezri, R., Holland, P., Schoborg, T. A., Abramovich, I., Takáts, S., Dillard, C., Jain, A., O’Farrell, F., Schultz, S. W., Hagopian, W. M., et al. (2021). Host autophagy mediates organ wasting and nutrient mobilization for tumor growth. EMBO J 40, e107336.

Kim, M.-J. and O’Connor, M. B. (2021). Drosophila Activin signaling promotes muscle growth through InR/TORC1-dependent and -independent processes. Development 148, dev190868.

Kwon, Y., Song, W., Droujinine, I. A., Hu, Y., Asara, J. M. and Perrimon, N. (2015). Systemic Organ Wasting Induced by Localized Expression of the Secreted Insulin/IGF Antagonist ImpL2. Developmental Cell 33, 36–46.

Lee, J., Ng, K. G.-L., Dombek, K. M., Eom, D. S. and Kwon, Y. V. (2021). Tumors overcome the action of the wasting factor ImpL2 by locally elevating Wnt/Wingless. Proc Natl Acad Sci U S A 118, e2020120118.

Leevers, S. J., Weinkove, D., MacDougall, L. K., Hafen, E. and Waterfield, M. D. (1996). The Drosophila phosphoinositide 3-kinase Dp110 promotes cell growth. The EMBO Journal 15, 6584–6594.

Lodge, W., Zavortink, M., Golenkina, S., Froldi, F., Dark, C., Cheung, S., Parker, B. L., Blazev, R., Bakopoulos, D., Christie, E. L., et al. (2021). Tumor-derived MMPs regulate cachexia in a Drosophila cancer model. Dev Cell S1534-5807(21)00638–9.

Manent, J., Banerjee, S., de Matos Simoes, R., Zoranovic, T., Mitsiades, C., Penninger, J. M., Simpson, K. J., Humbert, P. O. and Richardson, H. E. (2017). Autophagy suppresses Ras-driven epithelial tumourigenesis by limiting the accumulation of reactive oxygen species. Oncogene 36, 5576–5592.

Mirth, C., Truman, J. W. and Riddiford, L. M. (2005). The role of the prothoracic gland in determining critical weight for metamorphosis in Drosophila melanogaster. Curr Biol 15, 1796–1807.

Moraes, L. N., Fernandez, G. J., Vechetti-Júnior, I. J., Freire, P. P., Souza, R. W. A., Villacis, R. A. R., Rogatto, S. R., Reis, P. P., Dal-Pai-Silva, M. and Carvalho, R. F. (2017). Integration of miRNA and mRNA expression profiles reveals microRNA-regulated networks during muscle wasting in cardiac cachexia. Sci Rep 7, 6998.

Newton, H., Wang, Y.-F., Camplese, L., Mokochinski, J. B., Kramer, H. B., Brown, A. E. X., Fets, L. and Hirabayashi, S. (2020). Systemic muscle wasting and coordinated tumour response drive tumourigenesis. Nature Communications 11, 4653.

Ni, X., Yang, J. and Li, M. (2012). Imaging-guided curative surgical resection of pancreatic cancer in a xenograft mouse model. Cancer Lett 324, 179–185.

Parker, J. and Struhl, G. (2015). Scaling the Drosophila Wing: TOR-Dependent Target Gene Access by the Hippo Pathway Transducer Yorkie. PLoS Biol 13, e1002274.

Parkin, C. A. and Burnet, B. (1986). Growth arrest of Drosophila melanogaster on erg-2 and erg-6 sterol mutant strains of Saccharomyces cerevisiae. Journal of Insect Physiology 32, 463–471.

Pastor-Pareja, J. C. and Xu, T. (2011). Shaping Cells and Organs in Drosophila by Opposing Roles of Fat Body-Secreted Collagen IV and Perlecan. Dev Cell 21, 245–256.

Peterson, A. J., Jensen, P. A., Shimell, M., Stefancsik, R., Wijayatonge, R., Herder, R., Raftery, L. A. and O’Connor, M. B. (2012). R-Smad Competition Controls Activin Receptor Output in Drosophila. PLOS ONE 7, e36548.

Portela, M., Casas-Tinto, S., Rhiner, C., López-Gay, J. M., Domínguez, O., Soldini, D. and Moreno, E. (2010). Drosophila SPARC is a self-protective signal expressed by loser cells during cell competition. Dev Cell 19, 562–573.

Santabárbara-Ruiz, P. and Léopold, P. (2021). An Oatp transporter-mediated steroid sink promotes tumor-induced cachexia in Drosophila. Dev Cell 56, 2741–2751.e7.

Semaniuk, U., Piskovatska, V., Strilbytska, O., Strutynska, T., Burdyliuk, N., Vaiserman, A., Bubalo, V., Storey, K. B. and Lushchak, O. (2021). Drosophila insulin-like peptides: from expression to functions – a review. Entomologia Experimentalis et Applicata 169, 195–208.

Shahab, J., Baratta, C., Scuric, B., Godt, D., Venken, K. J. T. and Ringuette, M. J. (2015). Loss of SPARC dysregulates basal lamina assembly to disrupt larval fat body homeostasis in Drosophila melanogaster. Dev Dyn 244, 540–552.

Song, W., Kir, S., Hong, S., Hu, Y., Wang, X., Binari, R., Tang, H.-W., Chung, V., Banks, A. S., Spiegelman, B., et al. (2019). Tumor-Derived Ligands Trigger Tumor Growth and Host Wasting via Differential MEK Activation. Developmental Cell 48, 1–17.

Tiebe, M., Lutz, M., De La Garza, A., Buechling, T., Boutros, M. and Teleman, A. A. (2015). REPTOR and REPTOR-BP Regulate Organismal Metabolism and Transcription Downstream of TORC1. Dev Cell 33, 272–284.

Urrutia, H., Aleman, A. and Eivers, E. (2016). Drosophila Dullard functions as a Mad phosphatase to terminate BMP signaling. Sci Rep 6, 32269.

Weinkove, D., Neufeld, T. P., Twardzik, T., Waterfield, M. D. and Leevers, S. J. (1999). Regulation of imaginal disc cell size, cell number and organ size by Drosophila class I(A) phosphoinositide 3-kinase and its adaptor. 9, 1019–1029.

Yoo, S., Nair, S., Kim, H., Kim, Y., Lee, C., Lee, G. and Park, J. H. (2020). Knock-in mutations of scarecrow, a Drosophila homolog of mammalian Nkx2.1, reveal a novel function required for development of the optic lobe in Drosophila melanogaster. Developmental Biology 461, 145–159.

Zang, Y., Wan, M., Liu, M., Ke, H., Ma, S., Liu, L.-P., Ni, J.-Q. and Carlos Pastor-Pareja, J. (2015). Plasma membrane overgrowth causes fibrotic collagen accumulation and immune activation in Drosophila adipocytes. eLife 4, e07187.

Zhang, H., Stallock, J. P., Ng, J. C., Reinhard, C. and Neufeld, T. P. (2000). Regulation of cellular growth by the Drosophila target of rapamycin dTOR. Genes & Development 14, 2712–2724.

Zhang, Y., Billington, C. J., Pan, D. and Neufeld, T. P. (2006). Drosophila Target of Rapamycin Kinase Functions as a Multimer. Genetics 172, 355–362.

